# Image-based phenotypic sorting of synthetic cells

**DOI:** 10.1101/2025.09.25.678515

**Authors:** Marijn van den Brink, Marlena Stam, Nico J. Claassens, Christophe Danelon

## Abstract

Understanding the relationships between genotype and phenotype is key to many areas of biological research and to the development of synthetic cells. We describe an image-based screening and sorting workflow that explores the phenotypes of gene-expressing vesicles within nonclonal populations and selects the desired variants. Using automated confocal microscopy and real-time, neural-network-assisted image analysis, we demonstrate that liposomes can be selected for fluorescence intensity, protein localization, membrane morphology, and dynamic behaviors, and their phenotype can be linked to genetic content. This approach could substantially accelerate the evolution of cellular functions in a minimal synthetic context.

## INTRODUCTION

Modular reconstitution of basic cellular functions in liposomes has laid the groundwork for building up an entire synthetic cell. For instance, DNA replication^1^, transcription-translation^2–4^, phospholipid synthesis^5–8^, and membrane remodeling^9–14^ have already been reconstituted in liposomes. The functional states of the engineered liposomes are commonly assessed by fluorescence microscopy techniques, including imaging flow cytometry^15^. Image-based methods allow for the exploration of a large observation space, defined as the set of features making up a liposome’s phenotype. Therefore, vesicles exhibiting complex phenotypes, e.g., dynamic behaviors, spatial organizations and membrane morphologies, can be identified, expanding the range of target functions toward which liposomes can be engineered, ultimately accessing a high degree of “aliveness”.

When the liposome’s phenotype is directed by a gene expression program, methods of in vitro evolution can be applied to accelerate the search for a desired function^16,17^. Here, diversity is created by producing genetic variants directly inside liposomes^18^ or externally prior to random encapsulation^19^, leading to heterogenous—nonclonal—populations of liposomes exhibiting various, potentially distinct phenotypes. A major challenge resides in linking the specific phenotypes to their respective genotypes for sequence-function mapping. Pooled genetic screening of library variants in liposomes and isolation of the best-performing ones can be achieved by fluorescence-activated cell sorting (FACS)^19–25^. However, phenotypic interrogation here relies on unidimensional parameters, which restricts selection to simple properties.

Technological advances in image-based screening and sorting of cells are revolutionizing our understanding of genotype-to-phenotype relationships with great implications in single-cell omics and clinical research^26^. High-throughput image-activated cell sorters make sort decisions within micro-to milliseconds by real-time image analysis using predefined image parameters^27,28^ and/or more powerful machine learning strategies^29^ to classify complex phenotypes. However, drawbacks of these sorters are the low equipment accessibility, the limited spatial resolution, the limited number of fluorescence imaging channels, the inability to acquire time-lapse images or images in 3D (at multiple z planes), or the possible impact of the flow on some phenotypic traits^30^. Improved spatial resolution is obtained with imaging flow cytometry (IFC), but this technology lacks the ability to sort^31^. An alternative method that offers improved spatial and temporal resolution is the two-step combination of microscopy and FACS. Microscopy images are analyzed in real-time, whereafter the cells of interest are optically tagged for sorting by FACS. While this technique has been established for the analysis of mammalian and yeast cells^32–38^, its potential in the research field of synthetic cells remains unexplored.

In this study, we describe a confocal microscopy-based method that assesses the phenotypes of gene-expressing liposomes and marks variants that exhibit the desired properties through selective photoactivation. Photoactivated liposomes are then isolated by FACS, and the enrichment of the functional variants is determined by quantitative polymerase chain reaction (qPCR) or DNA sequencing. High-resolution image acquisition, analysis, and photoactivation are fully automated, employing trained neural networks and pre-defined parameters (e.g., intensity, size, circularity) for liposome classification. This approach allowed us to isolate liposomes with specific spatial and temporal features— including protein localization, membrane morphology and dynamic behaviors—from large populations of hundreds of thousands of liposomes per experiment. Our work thereby establishes microscopy-based phenotypic selection of gene-expressing liposomes as a core technology for the construction of synthetic cells using a system’s level evolutionary approach.

## RESULTS

### Establishing image-based liposome sorting

We aim to sort liposomes based on the visual phenotypes that arise from the expression of a library of different DNA variants. Synthesis of proteins that direct the phenotypic states is established directly in liposomes by the co-encapsulation of DNA and a cell-free transcription-translation system (PURE system)^39^ (**Fig. 1**). We used a laser-scanning confocal fluorescence microscope with a high numerical aperture objective to screen 50-150 liposomes per field-of-view (FOV) enabling precise examination of vesicle morphologies and subcellular features. The liposomes also encapsulate purified PAmCherry2, a photoactivatable mCherry, which marks individual liposomes for sorting when its fluorescence is activated by point-stimulation with 405-nm light. Automated image acquisition, multimodal image analysis, and stimulation of targeted liposomes iterate over each FOV, allowing for the screening of hundreds of thousands of liposomes per experiment of ∼8 hours. Next, the liposomes with photoactivated PAmCherry2 are isolated using FACS and the sorted DNA variants are analyzed by qPCR or PCR-amplified for DNA sequencing. The recovered variants could also be subjected to another round of diversification, re-encapsulation, and sorting.

**Fig. 1.**
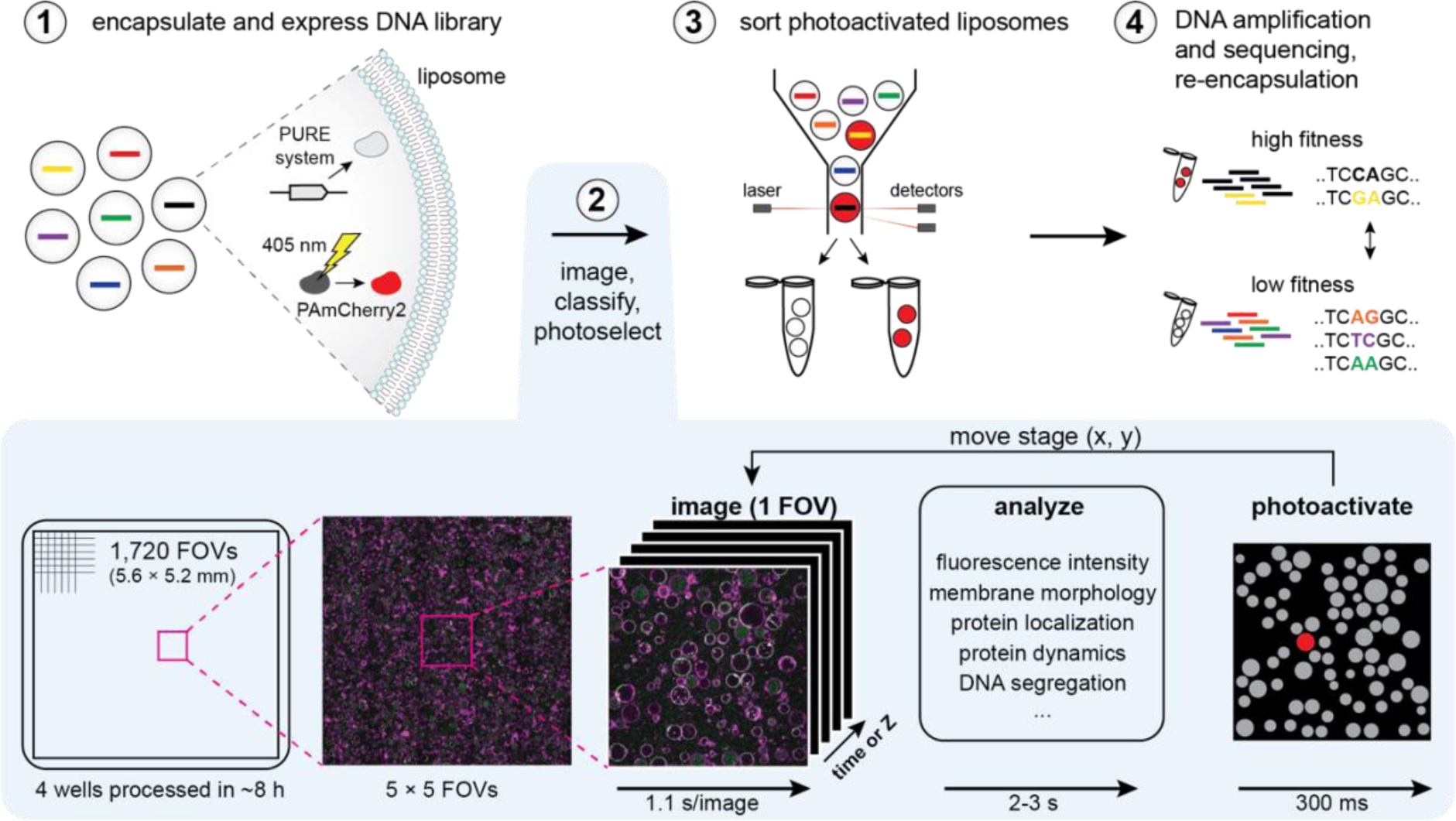
Schematic workflow of image-based liposome sorting. Liposomes encapsulate a DNA library alongside PURE system and purified PAmCherry2. Automated imaging, real-time analysis, and point-stimulation by 405-nm light to photoactivate PAmCherry2 in liposomes with a desired visual phenotype, is iterated for each FOV. The duration of each step is indicated. The photoactivated liposomes (in red on the right-most picture) are then sorted by FACS and DNA is recovered for analysis or re-encapsulation.

First, we verified that co-encapsulation of 20 µM purified PAmCherry2 was compatible with the expression of a DNA template encoding YFP (**Fig. 2a**). Point-stimulation (405-nm light at 84 µW laser power for 300 ms) of single liposomes located in the center xy coordinate of the FOV led to an increase in PAmCherry2 intensity in the entire lumen. Imaging at different z planes confirmed that, while a variety of liposome sizes was present (from 2 up to 20 µm in diameter) most of the vesicles were captured in a single z plane ∼2-3 µm above the glass surface (**Fig. 2a**). This is especially important for preventing off-target photoactivation in liposomes located within the cone-shaped illumination volume above and below the focal point. After photoactivating 29 liposomes with expressed YFP, all liposomes showed an increase in PAmCherry2 fluorescence intensity (**Fig. 2a**).

**Fig. 2.**
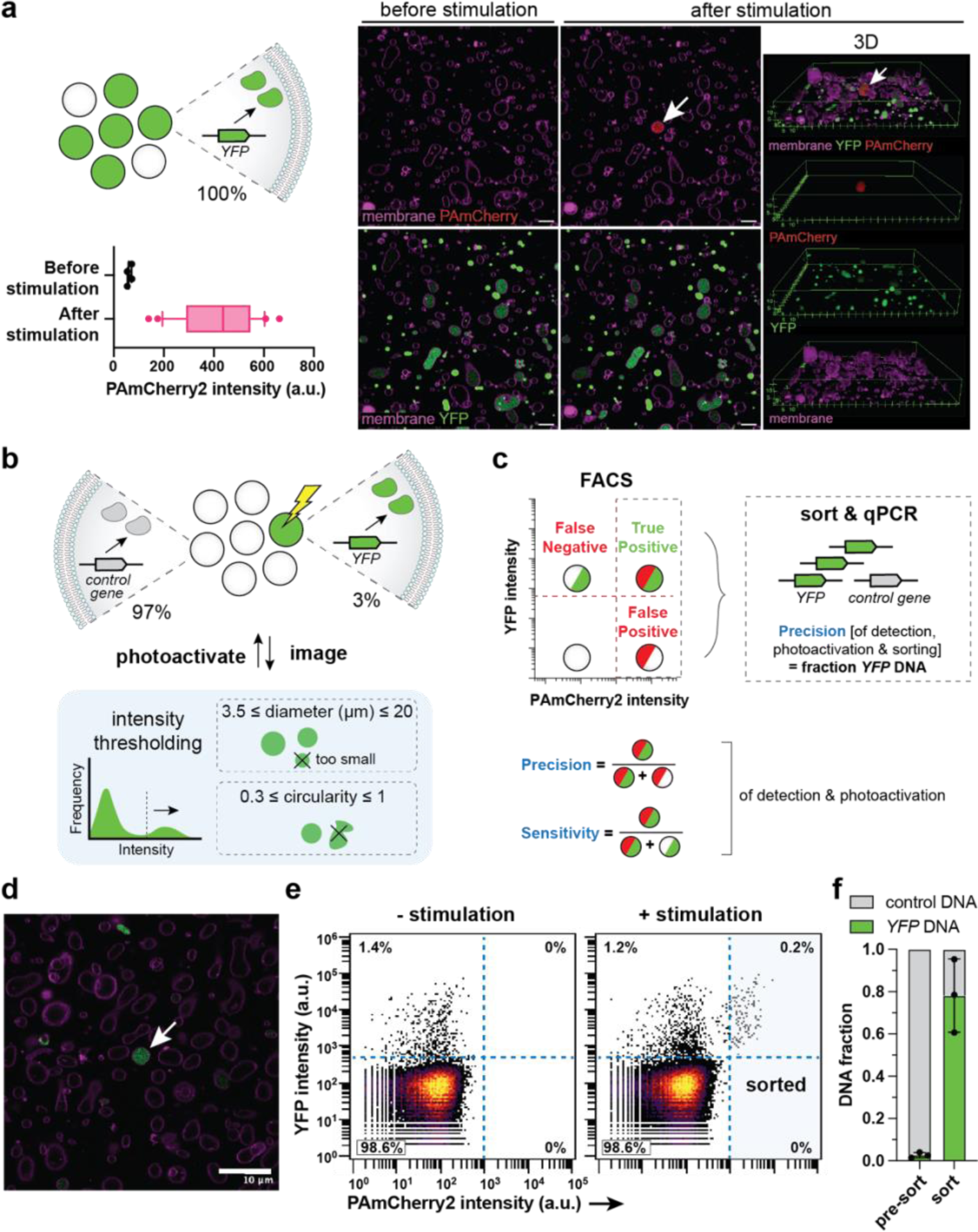
Image-based sorting of YFP-expressing liposomes. **a** Photoactivation of PAmCherry2 in YFP-expressing liposomes. Liposomes contained 5 nM of *YFP* DNA template and 20 µM of purified PAmCherry2. Stimulation was performed in the center of the FOV using 84 µW laser power for 300 ms. The PAmCherry2 fluorescence intensity was measured 10 seconds after stimulation. Graph displays n=29 liposomes. Scale bars: 10 µm. **b** Image-based sorting of YFP-expressing liposomes from a two-gene mock library prepared by mixing two populations of liposomes at a 1:39 volume ratio, each containing an equal amount of linear DNA template, either *YFP* or *btubB* (control gene). Automated image analysis was performed using intensity thresholding on the YFP signal. **c** The precision and sensitivity of the method were assessed using FACS and qPCR analysis of the *YFP* and control DNA in the sorted liposomes. **d** Example microscopy image of the mock library. Green, YFP; magenta, Cy5-conjugated lipids. Scale bar: 10 µm. Arrow indicates liposome selected for photostimulation (300 ms at 34 µW (replicates 1 and 2) or 84 µW (replicate 3) laser power). **e** Flow cytometry measurements of liposome populations without (28,898 events) or with (54,490 events) stimulation. **f** Fractions of *YFP* and control DNA in the sorted and pre-sort liposome samples. n=3 biological replicates (distinct mixed liposome populations).

Next, we examined the accuracy of PAmCherry2 photoactivation for different laser powers (34, 84 or 167 µW). All target liposomes were successfully tagged for the three photoactivation conditions with no significant differences in the median intensity values (**Fig. S1a**). We then measured the lumen intensities of all other liposomes within 15-µm xy distance to the point of stimulation and found activation precisions of 96% at 34 µW, 89% at 84 µW, and 86% at 167 µW (**Fig. S1a**). Thus, 34 µW laser power is the preferred choice, enabling photoactivation of all target liposomes, while limiting off-target activation.

Interestingly, we observed a positive correlation between the fluorescence intensities of YFP and activated PAmCherry2 (**Fig. S1b**). Specifically, liposomes that did not show YFP expression, did also not exhibit an increase in PAmCherry2 signal after photostimulation. This result suggests that encapsulation of macromolecules during liposome formation may be an all-or-nothing (or poorly efficient) process, whereby most components in the swelling solution would either be entrapped together or not all. Furthermore, the YFP signal did not bleach upon photostimulation (**Fig. S1c**), and the activated PAmCherry2 was stable for at least 21 hours, giving enough time to sort the tagged liposomes by FACS (**Fig. S1d-e**).

### Image-based enrichment of YFP-expressing vesicles

To assess the performance of the imaged-based selection pipeline, we quantified the precision and sensitivity for sorting YFP-expressing liposomes starting from a two-gene mock library. A mixed population of liposomes containing PAmCherry2, PURE system, and either a *YFP* gene or a control gene (1:39 volume ratio) was processed (**Fig. 2b-d**). The YFP-expressing liposomes were identified in real-time using basic analysis parameters including intensity thresholding, object size, and circularity, and were photoactivated. Per replicate experiment, we screened liposomes in 2 to 5 thousand FOVs (31 to 77 mm^2^), and photoactivated ∼800 to 7,200 liposomes (**Table S1**). Stimulated and non-stimulated samples were analyzed by flow cytometry (**Fig. 2e**, **Fig. S2**, **Table S2**). Approximately 1 to 2% of the liposomes were expressing YFP. This percentage is in agreement with the ratio at which the liposome populations were mixed (1:39) and the percentage of YFP-expressing liposomes in the initial population prior mixing with the control population (40 to 80%). While the non-stimulated sample only showed liposomes with a low PAmCherry2 signal, the stimulated sample yielded 0.3% ± 0.1% of the liposomes with a high PAmCherry2 signal (n=3 biological repeats). By processing the events in the four quadrants as indicated in **Fig. 2c**, we found that the YFP-expressing liposomes were detected and photoactivated with a precision of 100% (using laser power 34 µW) (**Fig. 2e**). Two replicate experiments with laser power 84 µW and where stimulation was not restricted to center of FOV resulted in a precision of 47% ± 8% (**Fig. S2**). Additionally, 15.3% ± 0.2% (n=3) of the YFP-expressing liposomes were photoactivated. This value defines the sensitivity of the method but is dependent on the stringency of the liposome classification. Ten to 20% of the photoactivated liposomes were detected by FACS and sorted, corresponding to 150 to 900 liposomes (**Table S1**). Lastly, the enrichment of the *YFP* gene over the control gene was quantified by qPCR (**Fig. 2f**, **Table S3**). The percentage of *YFP* amount increased from 3% ± 1% to 78% ± 17%, corresponding to a 39 ± 28 fold enrichment (n=3).

We further investigated the possible contribution of off-target stimulation to the presence of the control gene in the sorted fraction by directly sorting YFP-expressing liposomes from the mixed population by FACS, i.e., without imaging. Here, we obtained a percentage of *YFP* in the DNA pool of 81% ± 6% (n=4) after sorting (**Fig. S3**, **Tables S4-S6**) (to be compared to 78% with the image-based workflow), indicating that photostimulation did not significantly affect the sorting precision.

### Real-time, neural network-assisted detection of liposomes and spatial protein organization

Deep learning models have become state-of-the-art for image analysis thanks to their high prediction speed, robustness to biological variability, noisy data and varying imaging conditions, and reduced bias from manual parameter tuning^40,41^. Therefore, the single-stage object detector You Only Look Once (YOLO)^42–44^ was chosen and used to identify liposomes in real-time based on their membrane morphology and the spatial organization of fluorescent proteins. As a proof-of-principle, we applied YOLO to detect proteins of the Min system, a regulator of bacterial cytokinesis and a candidate for the reconstitution of division processes in synthetic cells^45,10^. We expressed the membrane binding protein MinD from a *minD* DNA template and added the purified eGFP-MinC fusion protein as a reporter of the presence of active MinD (**Fig. 3a**). The localization of eGFP-MinC in the lumen would indicate the absence of MinD, while its localization at the membrane would reveal the functional expression of MinD.

**Fig. 3.**
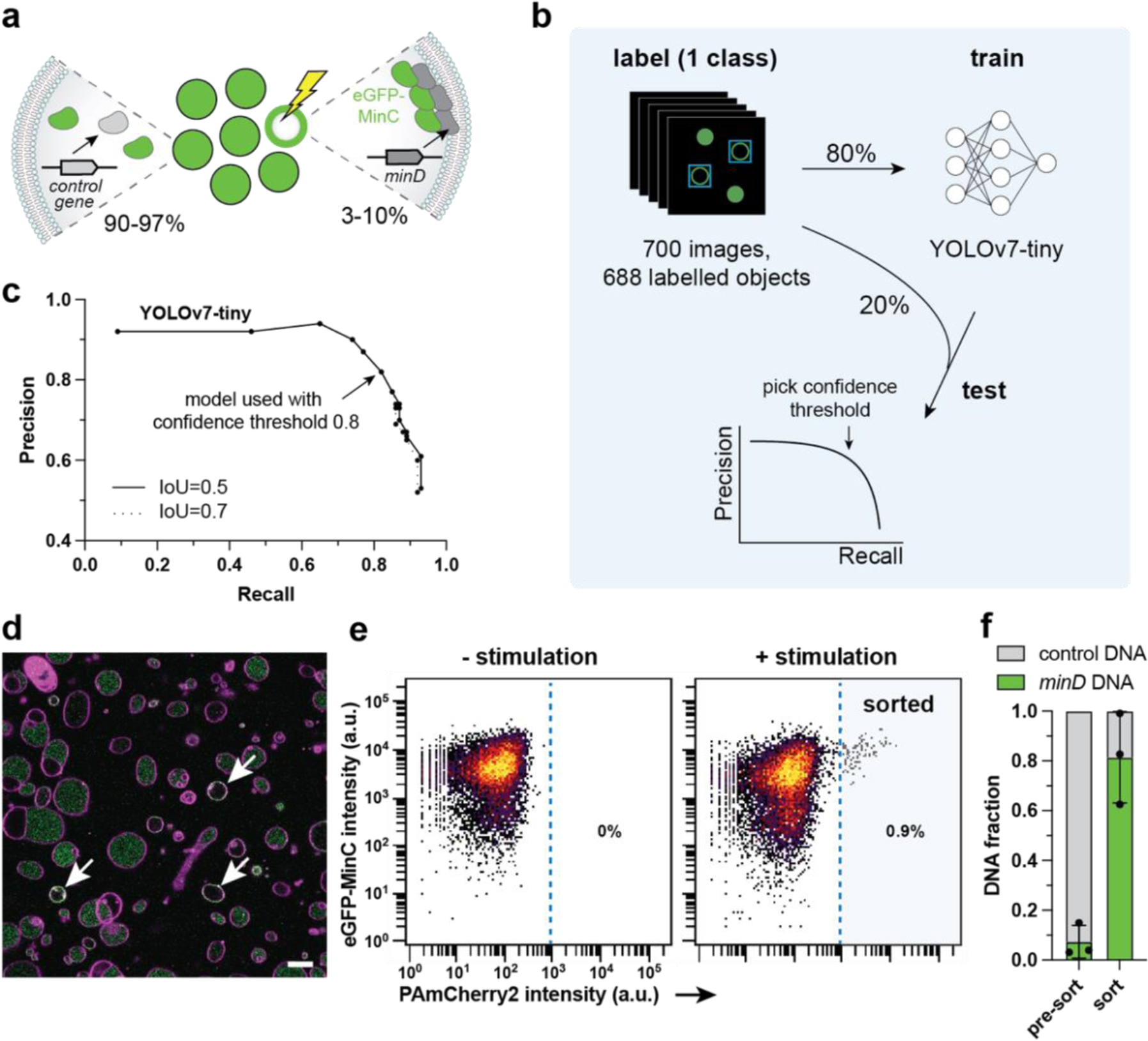
Image-assisted sorting of membrane-localized MinD using deep learning. **a** A mock DNA library was prepared by mixing two populations of liposomes (1:9 or 1:39 volume ratio) expressing either *minD* or *btubB* (control gene) and containing 500 nM of purified eGFP-MinC. When MinD is expressed, it binds to the membrane and recruits eGFP-MinC. **b** Labelling, training, and testing of YOLOv7-tiny using images displaying the membrane reporter dye Cy5 and GFP signals. **c** Performance of the trained YOLOv7-tiny model. IoU is Intersection over Union. **d** Example microscopy image of the mock library. Green, eGFP-MinC; magenta, Cy5-conjugated lipids. Scale bar: 10 µm. Automated image analysis was performed by the trained YOLOv7-tiny model. Arrows indicate liposomes classified for photostimulation during 300 ms at 34 µW or 84 µW laser power (Table S7). **e** Flow cytometry measurements of liposome populations without (10,365 events) or with (10,262 events) stimulation. Between 170 and 900 liposomes were sorted per replicate. **f** Fractions of *minD* and control DNA in the sorted and pre-sort liposome samples. n=3 biological replicates.

As training data, we used 700 images of liposomes containing both the lipid membrane and eGFP-MinC channels (**Fig. 3b**). The dataset originated from multiple liposome samples (biological replicates) to increase the model’s robustness to variation in sample preparation and imaging conditions. Each image displayed many liposomes with various phenotypes: no eGFP-MinC signal, eGFP-MinC on the membrane or/and in the lumen, unilamellar or multilamellar vesicles, circular liposomes or structures with other shapes, and a wide distribution of sizes. We annotated 688 objects in one class, aiming for unilamellar, circular liposomes of 3 to 15 µm in diameter with eGFP-MinC on the membrane. Using 80% of the labelling dataset, we trained YOLOv3-tiny^43^ with pretrained weights, and both YOLOv3-tiny and YOLOv7-tiny^44^ with and without pretrained weights (**Fig. S4a**). We tested the two models using the remaining 20% images and found that the best-performing model was YOLOv7-tiny with a precision of 82% and recall of 82% at a confidence threshold of 0.8 (**Fig. 3c**). Communication between the image acquisition software and the trained YOLOv7-tiny model was established for real-time image analysis and photoactivation.

Two populations of liposomes containing PAmCherry2, PURE system, purified eGFP-MinC, and either the *minD* gene or a control gene were mixed in a 1:9 (n=1) or 1:39 (n=2) ratio (*minD* : control gene) (**Fig. 3a**). Liposomes exhibiting eGFP-MinC signal at the membrane were photostimulated to select vesicles containing *minD* (**Fig. 3d**, **Table S7**). Photoactivated liposomes representing 0.03 to 0.94% of the total population were sorted by FACS (**Fig. 3e, Fig. S5, Table S8**), and the fractions of gene variants were quantified by qPCR in the pre-sort and sorted liposome samples. The percentage of *minD* increased from 7% ± 7% in the pre-sort sample to 81% ± 18% after sorting, corresponding to a 19 ± 12 fold enrichment (n=3 biological repeats) (**Fig. 3f, Table S9**). Because neural network analysis can be applied to identify images with rich spatial and morphological information, these results show the potential of our method for the selection of synthetic cells with complex phenotypes from heterogeneous samples.

### Video-based photoactivation of liposomes displaying dynamic protein oscillations

As flow-based image-activated sorters can only detect static features, we sought to exploit our method to photoactivate liposomes based on dynamic processes. In *E. coli*, the Min system undergoes oscillatory behaviors that aid in positioning the division site^46^. In liposomes, co-expression of MinD and MinE leads to different types of dynamic behavior, including pulsing, circling, and pole-to-pole oscillations with periods typically ranging from 15 to 45 seconds (**Fig. 4a**)^10^. Liposomes can also exhibit an uncategorized or halted dynamic mode, and modes can switch from one to another or change directionality, leading to a large variability in dynamic and static phenotypes within one liposome sample. Consequently, time-lapse microscopy is required to assess whether a liposome exhibits a dynamic or static organization of Min proteins. To detect the temporal behaviors of the Min proteins in real time, the liposomes were imaged with 12 second intervals over a period of 48 seconds. We trained YOLOv7-tiny to detect two classes: class 1 consists of liposomes with eGFP-MinC not located on the membrane, while class 2 consists of liposomes with eGFP-MinC (partly) located on the membrane (**Fig. 4b**). The model was trained on 80% of the images and tested using the separate 20% of the dataset, yielding a precision of 83% and recall of 86% at a confidence threshold of 0.9 (**Fig. S4b**). Hereafter, we retrained a model on the complete labelled dataset (700 to 900 objects per class).

**Fig. 4.**
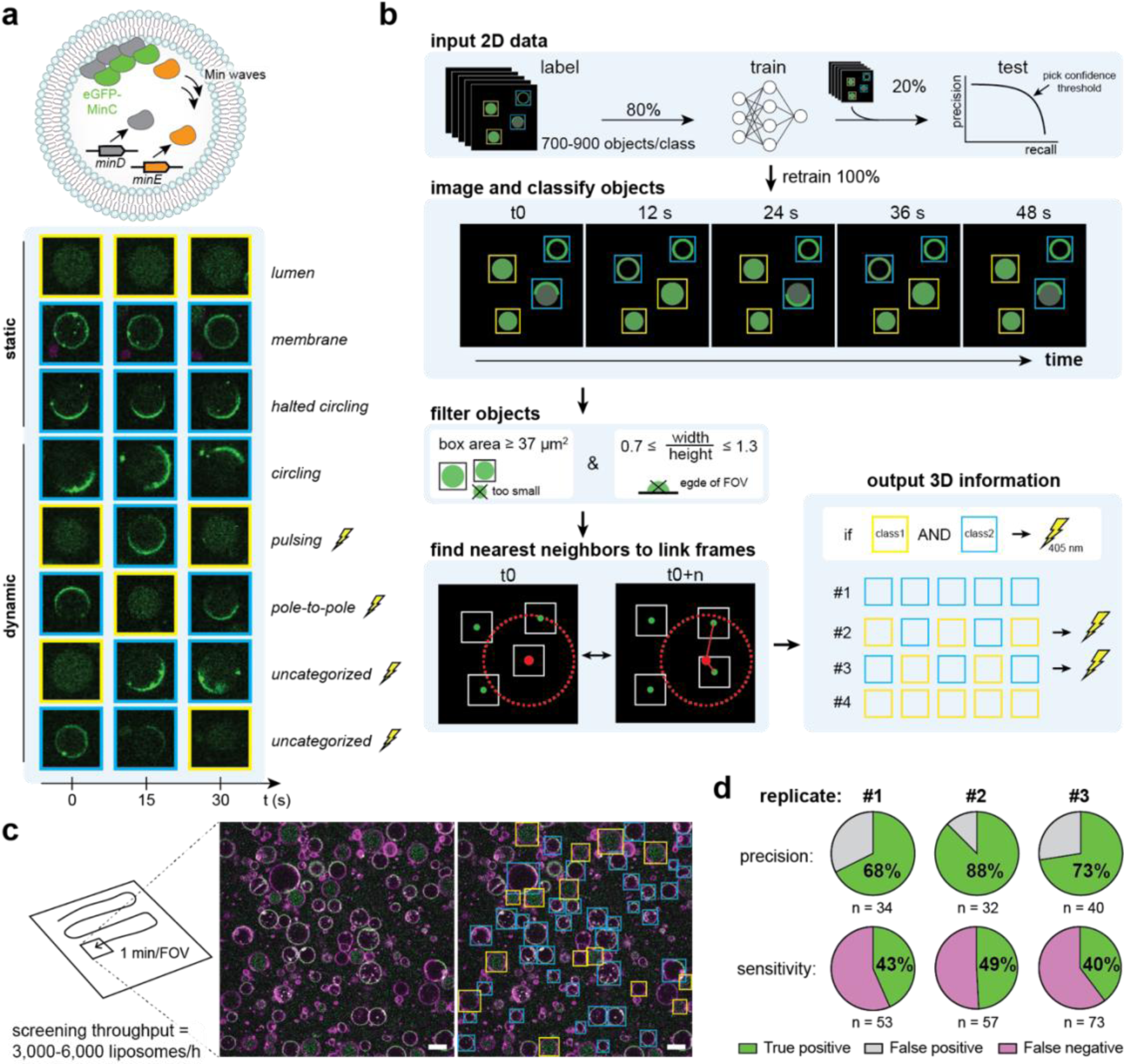
Real-time detection and tagging of liposomes with dynamic Min oscillations. **a** Min oscillations driven by the co-expression of MinD and MinE were visualized using purified eGFP-MinC. Representative images of eight liposomes exhibiting static or dynamic behaviors. Only liposomes with dynamic patterns were photoactivated. **b** Automated image analysis pipeline. Liposomes were detected using a YOLOv7-tiny model, trained on 2D data using two classes (indicated with blue and yellow bounding boxes). To assure accurate liposome tracking across time frames, small liposomes (diameter < 6 µm), which tend to move in and out of focus more than larger liposomes, and liposomes with aspect ratio below 0.7 and above 1.3, were excluded. Linking multiple time frames through a nearest neighbor search enabled the discrimination of static vs dynamic behaviors. **c** Automated acquisition of liposome images and real-time identification of static vs dynamic self-organization of Min proteins. Scale bar: 10 µm. **d** Performance of the classification pipeline. Precision and sensitivity were calculated relative to the ground truth, which entailed manual classification of the liposomes in the videos. n=3 biological replicates. Stimulation time was 300 ms, and laser power was 34 µW. The YOLOv7-tiny model from replicate 1 was retrained with 9 additional images to strengthen the model (Fig. S4b). For replicates 1 and 2, a population of liposomes expressing MinD and MinE was mixed 1:1 with liposomes expressing MinD and a control protein (BtubB). For replicate 3, only liposomes expressing MinD and MinE were used.

Detected liposomes were tracked across multiple time frames using a nearest-neighbor search (**Fig. 4b**). While most liposomes remained stationary during the measurements, a few moved substantially in xy direction, increasing the likelihood of being misidentified as a different liposome. Therefore, we applied a 5-µm maximum displacement threshold relative to the position in the first frame. Then, each class for a given liposome was compared over the consecutive time frames. The change from one class to the other reveals a dynamic event and these liposomes were photoactivated. In 8 hours, we screened approximately 40,000 liposomes and stimulated one liposome in every 2 FOVs (∼0.5% of all liposomes) (**Fig. 4c**, **Table S10**). Manual labelling of liposomes showing both the presence and absence of membrane-localized eGFP-MinC across different time frames resulted in a precision of 76 ± 10% and a sensitivity of 44 ± 5% (n=3 biological repeats), validating video-based classification of liposomes (**Fig. 4d**).

### Selection of best-performing membrane-binders from a library of *minD* variants

At last, we showed that image-based screening and sorting can be successfully coupled to the recovery and sequencing of DNA from the sorted liposomes. We sought to find the best-performing variants within a *minD* library harboring mutations in the hydrophobic patch of the membrane targeting sequence (MTS)^47^, most likely conferring different abilities to bind to the liposome membrane. Nine DNA variants were separately encapsulated and expressed in liposomes, whereafter the clonal liposome populations were mixed and subjected to the image-based selection pipeline to sort the liposomes with membrane-localized eGFP-MinC (**Fig. 5b**, **Fig. S6**, **Tables S11-S12**). The DNA was recovered by PCR from 88 up to 704 sorted liposomes per biological replicate (**Fig. 5c**, **Fig. S7c**), and the fraction of each variant before and after sorting was quantified by Nanopore sequencing (**Fig. 5d**). From the nine variants present in the input library, only two were found in the sorted pool: Var1 (the wild type) and Var7 (amino acids: FL>LH). Encapsulation and expression of the individual variants confirmed that Var1 and Var7 encode a membrane-binding MinD, while the other mutants could not (**Fig. 5e, Fig. S7a-b**). These results are in agreement with a previous study that showed that substitution of hydrophobic amino acids to charged amino acids in the MTS caused defective binding of MinD to the membrane in *E. coli*^47^. Altogether, these findings demonstrate the feasibility to isolate and genotype active gene variants from a library using image-based liposome phenotyping.

**Fig. 5.**
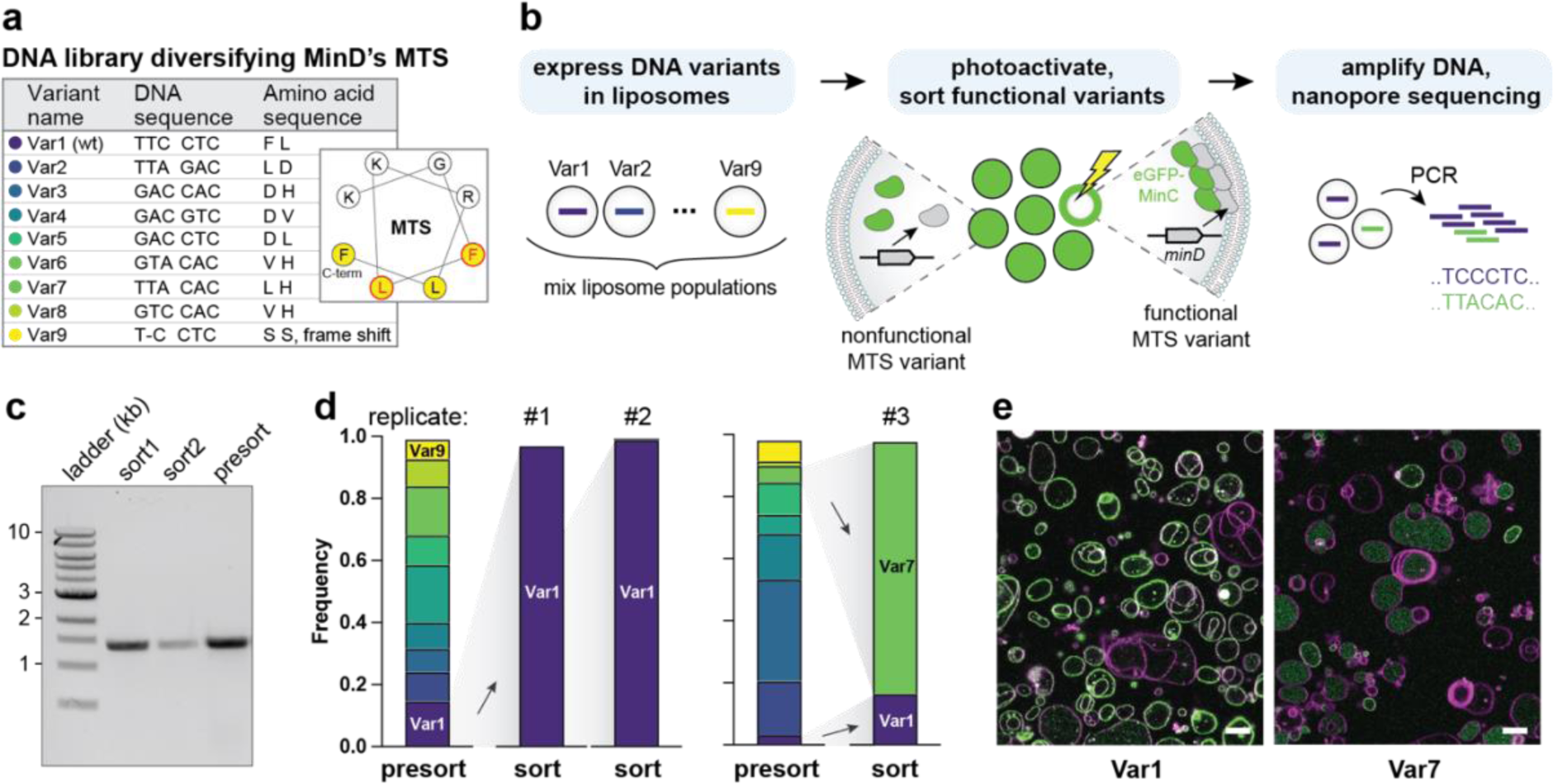
Enrichment of active *minD* variants from a library. **a** Design of the library of MTSs in the gene *minD*. Mutations were introduced in two of the codons in the hydrophobic patch of the highly conserved MTS motif of MinD. **b** Image-based sorting of liposomes expressing MinD variants with different affinities to the membrane. Populations of clonal liposomes, each containing one library variant, were pooled and subjected to image-based selection. Mutations lead to differential localization of eGFP-MinC to the membrane (functional MTS) or to the lumen (nonfunctional MTS). DNA was amplified from the sorted liposomes by PCR and sequenced. Stimulation time was 300 ms and laser power was 34 µW. **c** Gel electrophoresis with PCR-amplified *minD* (1,398 bp) recovered from liposomes sorted by FACS. **d** Frequency of the DNA variants was quantified using Nanopore sequencing before and after liposome sorting. Two starting ratios of the nine DNA variants were prepared, from which in total three sorting replicates were performed. **e** Confocal microscopy images of the two enriched variants (Var1 and Var7) separately encapsulated and expressed in liposomes. Green, eGFP-MinC; magenta, Cy5-conjugated lipids. Scale bar: 10 µm.

## DISCUSSION

We describe a confocal microscopy-based method that explores the phenotypes of gene-expressing liposomes within large populations and sorts the desired variants using photo-tagging followed by FACS. Our results offer a new promising way to accelerate the development of synthetic cells through in vitro evolution whereby the system’s fitness is assessed by visual screening. With this objective in view, we here discuss the current assets of the proposed technology compared to existing ones, as well as possible extensions that could enable the discovery of phenotypically complex synthetic cells from a large library of genetic and molecular variants.

Unlike traditional methods that enrich variants based on one-dimensional readouts in time and space, such as FACS^19–25^, affinity-based selection^17,48^, and self-selection through DNA replication^18^, visual phenotyping allowed us to classify liposomes based on morphology and spatiotemporal protein organization. While photoselection-based methods similar to ours have been developed for biological cells^32–38^, working with liposomal synthetic cells imposes several challenges: the small size (diameter < 10 µm), polydispersity and low phase-contrast of the vesicles, stochastic encapsulation of the photoactivatable probe, and positioning of the liposomes on a 2D surface. Because of these constraints, we chose confocal-fluorescence imaging using a high numerical aperture (NA 1.49, 100× oil-immersion) objective to measure with a high spatial resolution. Despite the small size of the liposomes, photoactivation of single vesicles was accurate and sensitive; all liposomes that displayed gene expression also exhibited higher PAmCherry2 fluorescence after stimulation (**Fig. 2, Fig. S1**).

In this work, we explored a combined set of liposome features including fluorescence intensity, protein co-localization to the membrane or lumen, and liposome morphology. We also introduced a temporal dimension by linking these phenotypes across time frames. By capturing 50 to 150 liposomes per FOV (128 µm × 128 µm), we screened up to 700,000 liposomes per experiment of ∼8 hours (including photoactivation). We demonstrated successful recovery and sequencing of the DNA underlying the selected phenotypes for typical sorting fractions of 150 to 1,000 liposomes, and we could even recover the DNA from as low as 88 sorted liposomes. We showcased the enrichment of the most active DNA variants from a library of nine variants (**Fig. 5**). The next step in image-based sorting of synthetic cells is enlarging the library size to make better use of the available screening capacity—e.g., a library of 14,000 variants that will be screened with 50-fold coverage. This will require establishing a method for multiplexing liposome production, where many liposome samples can be prepared in parallel, for example using lab automation. Alternatively, the DNA library could be encapsulated in a single liposome sample at a low DNA concentration (∼1 copy per liposome), but this strategy demands an increase in the fraction of ‘active’ liposomes to enable screening of large libraries, for instance by coupling gene expression with clonal amplification^49^. Such approaches should go hand-in-hand with smart library design and active learning-assisted directed evolution to enable efficient screening of large genetic and molecular spaces by focusing experimental testing to a smaller subset containing the most informative variants^50,51^.

While the screening throughput depends on the number of stimulated liposomes per FOV, it is on par with that of existing methods applied to biological cells. Yet it is low compared to flow-based imaging sorters that can typically analyze samples at a speed of 10^2^ to 10^3^ events per second^27,29^. Strategies to increase the throughput include (combinations of) shortening (i) the acquisition time by using a spinning-disk confocal microscope, (ii) the analysis time by processing multiple FOVs simultaneously or speeding up the communication between acquisition and analysis, and (iii) the stimulation time while increasing the 405-nm laser power. Noteworthy, ultrawide-FOV microscopes could dramatically increase the throughput by imaging FOVs of ∼5 to 42 mm^2^ at millisecond temporal resolution^36,37,52^. However, a better tradeoff will have to be found for improving the spatial resolution (NA objective 0.25 or 0.5^36,37,52^) as it is currently insufficient for the characterization of internal liposome structures. The same limitation of the spatial resolution also applies to the flow-based imaging sorters IACS2.0^29^ and BD FACSDiscover S8^27,28^.

We observed that only ∼10% of the photoactivated liposomes were commonly detected by FACS (**Tables S1, S7, S11**), which could limit the efficiency of the sorting method when the phenotypes of interest are extremely rare. This percentage is somewhat lower than the recovery rates reported for cells^33,38^. It is likely that the disruption of liposomes, incomplete harvesting during transfer from the imaging well to the FACS instrument, or the low scattered light intensity of the liposomes explain this observation.

High-content imaging, which captures multiple features of gene-expressing liposomes, enables multi-objective optimization that reflects the integrated activities of several biological processes. As the number of reconstituted biological modules increases, the liposome phenotypes become more complex, and the observation space becomes larger. Therefore, sorting multiple subpopulations would be very useful to obtain a more holistic view of the effects of genetic and molecular variations on the phenotype—beyond the best-performing variants only. This allows better identification of neutral and deleterious mutations and may provide deeper insights into mutational effects across different genetic or molecular backgrounds^53^. Sorting multiple subpopulations might be achieved by varying the activation time or laser power and sort multiple phenotypes by FACS based on PAmCherry2 intensity thresholding^32,34^. However, the overlap of PAmCherry2 intensity distributions at laser powers spanning 34 to 167 µW currently challenges multi-way sorting of photoactivated liposomes. Adjusting both the laser power and stimulation time might generate distinct PAmCherry2 intensity profiles.

In addition to DNA sequencing, the sorted liposome fractions could be subjected to other analytical investigations to reveal a rich array of cellular properties and functions, which can be coupled to the fitness level and genotype of each subpopulation. Recent advances in single-cell ‘omics’ could be applied to gene-expressing liposomes for mapping genotype and detailed molecular information, e.g., proteins, RNA transcripts and metabolites^36,54,55^.

Lastly, the current classification strategy relies on pre-defined metrics of physical parameters or on supervised machine learning with liposome images annotated by the user. This approach introduces potential human bias that narrows the exploration of phenotypic states. Unsupervised methods based on the structure of the data itself and lacking an explicit optimization target may increase robustness and expand the range of possible discoveries by revealing difficult-to-identify or hidden properties of the liposomes^56^. Implementing unsupervised learning to our imaging-based pipeline would lay the foundation for open-ended evolution of synthetic cells^16^.

## METHODS

### Reagents and plasmid DNA

Solutions were prepared using Milli-Q water with 18.2 MΩ resistivity (Merck Millipore). Reagents were purchased from Sigma-Aldrich unless indicated otherwise. Primers and plasmids are listed in **Tables S13** and **S14**, respectively. Plasmids G365 (pUC-*YFP*) and G437 (pUC-*minD*) were constructed as described earlier^49^. G439 (pUC-*btubB*) was constructed as described earlier^12^. G376 (pUC-*minE_opt*) was ordered from GenScript, as described earlier^10^. The expression vector G595 (pET15b-*PAmCherry2*) was constructed by cloning of the *PAmCherry2.1* gene^57^ (amplified with primers 1429 ChD and 1430 ChD) into the pET15b backbone (amplified with primers 1431 ChD and 1432 ChD) using Gibson Assembly. The expression vector G577 (pET15b-mEos2) was constructed by cloning of the *mEos2* gene (amplified from G571 with primers 1355 ChD and 1356 ChD) into the pET15b backbone (amplified with 1353 ChD and 1354 ChD) using Gibson Assembly.

### Protein purification

eGFP-MinC was purified as described in published protocols^58^. To synthesize PAmCherry2 and mEos2, 10 mL of overnight culture of *E. coli* BL21 (DE3) (New England Biolabs) carrying G595 (PAmCherry2) or G577 (mEos2) was diluted 1:100 in Lysogeny Broth (LB) medium containing 50 µg mL^−1^ ampicillin. The diluted culture was grown in a shaking incubator at 37 °C until the OD_600_ reached 0.4-0.6. Gene expression, under control of the T7-LacO promotor and in frame with an N-terminal His-tag, was induced by addition of 0.25 mM (PAmCherry2) or 1 mM (mEos2) isopropyl-β-D-thiogalactopyranoside (IPTG). The cells were incubated overnight at 16 °C in a shaking incubator (PAmCherry2) or at 26 °C for 4 h (mEos2). The cells from 1 L of culture were pelleted by centrifugation and resuspended in 10 mL of lysis buffer (50 mM HEPES-KOH pH 7.5, 500 mM NaCl and 10% glycerol). The cells were sonicated in an ice bath six times 20-30 seconds with an interval of 1 min and an amplitude of 40%. After centrifugation at 16,000 g for 15 min at 4 °C, the cell-free extract (supernatant) was harvested. 10 mM imidazole and SetIII protease inhibitor (EDTA-free, 1:1000 dilution, Calbiochem) were added. The cell-free extract was mixed with a HisPure Ni-NTA resin (Thermo Scientific), priorly equilibrated with lysis buffer, and incubated at 4 °C for 2 hours while tumbling. The unbound fraction was removed by spinning the resin at 750 g for 2 min. The resin with bound protein was washed twice with 20 equivalents (volume) of wash buffer (50 mM HEPES-KOH pH 7.5, 500 mM NaCl, 50 mM (PAmCherry2) or 40 mM (mEos2) imidazole, 10% glycerol). The protein was eluted with 6 mL of elution buffer (50 mM HEPES-KOH pH 7.5, 500 mM NaCl, 500 mM imidazole, 10% glycerol). The fluorescent fractions (for mEos2) were pooled together and buffer-exchanged with storage buffer (50 mM HEPES-KOH pH 7.5, 50 mM (PAmCherry2) or 100 mM (mEos2) NaCl, 10% glycerol) using a 10 kDa MWCO Amicon Ultra-15 centrifugal filter unit (Merck). Protein concentration was determined with a Bradford assay.

### Preparation of linear DNA templates for encapsulation in liposomes

Linear DNA templates for in-liposome expression were amplified by PCR from plasmids G365 (*YFP*), G439 (*btubB*), G437 (*minD*), G376 (*minE*) with primers 757 ChD and 709 ChD annealing to the T7 promoter and T7 terminator, respectively. The PCR reactions were set up at 50-µL volume and contained 200 µM of dNTPs, 500 nM of each primer, 200 pg µL^−1^ of plasmid DNA and 1 unit of Phusion High-Fidelity DNA Polymerase (Thermo Scientific) in 1× Phusion HF Buffer. The thermocycling reaction was performed as follows: 98 °C for 30 s, (98 °C for 10 s, 56 °C for 20 s, 72 °C for 30 s) × 30, and 72 °C for 10 min. The amplicons were purified directly or by gel extraction using the QIAquick PCR & Gel Cleanup Kit (Qiagen). The DNA concentration was measured with the Qubit dsDNA High Sensitivity Assay Kit and the Qubit 4 Fluorometer (Invitrogen).

### Synthesis of *minD* plasmid library and isolation of single variants

A 32-variant *minD* plasmid library (G598) with mutations in the MTS of MinD was synthesized by full-plasmid amplification of plasmid G437 (pUC19-*minD*) with a mutagenic primer set and subsequent re-circularization using the protocol adapted from Galka et al.^59^. PCR amplification was performed with primers 1445 ChD and 1448 ChD, of which the latter contains degeneracy at the MTS. The PCR reaction was set up at 20-µL volume and contained 400 µM of dNTPs, 300 nM of each primer, 350 pg µL^−1^ plasmid DNA and 0.4 units of KOD Xtreme Hot Start DNA Polymerase in 1× KOD Xtreme PCR Buffer and 5% DMSO. A touchdown thermocycling reaction was performed as follows: 94 °C for 2 min, (98 °C for 10 s, 74 °C for 4.2 min) × 4, (98 °C for 10 s, 72 °C for 4.2 min) × 4, (98 °C for 10 s, 70 °C for 4.2 min) × 4, (98 °C for 10 s, 60 °C for 20 s, 70 °C for 4.2 min) × 12, and 70 °C for 5 min.

Because the two primers are (partly) complementary, the two ends of the amplicon were identical, enabling re-circularization by mixing 18 µL of the PCR reaction with 54 µL of a Gibson reaction mix containing 1.3× ISO Buffer (0.5 M of Tris-HCl pH 7.5, 50 mM of MgCl_2_, 4 mM of dNTP mix, 50 mM of DTT, 0.25 g mL^−1^ of PEG-8000, 5 mM NAD), 0.053 U µL^−1^ of T5 exonuclease (Epicentre), 0.0033 U µL^−1^ of Phusion High-Fidelity DNA polymerase (Thermo Scientific), 5.3 U µL^−1^ of Taq DNA ligase (NEB) and 50 U mL^−1^ of DpnI endonuclease (NEB). The reaction was incubated in a thermocycler at 50 °C for 2 hours. The full reaction was loaded onto a 0.7% agarose gel, and the band corresponding to the circularized amplicon was purified by gel excision and further purified using QIAquick PCR & Gel Cleanup Kit (Qiagen).

Next, 50 µL of competent *E. coli* K-12 DH5α cells were transformed with 3 ng of purified plasmid using heat shock. Single colonies were picked from ampicillin-supplemented LB agar plates and grown overnight in ampicillin-supplemented (50 µg mL^−1^) LB medium. The plasmids were isolated with the PureYield Plasmid Miniprep System (Promega), and their sequences were analyzed by Sanger sequencing (Macrogen Europe, Netherlands) using primer 709 ChD.

### Liposome preparation

The swelling solutions (10 μL) were prepared on ice by mixing 5 μL of PURE*frex* 2.0 Solution I, 0.5 μL of Solution II, 1 μL of Solution III, 0.75 U μL^−1^ SUPERase·In RNase Inhibitor (Ambion), 20 μM purified PAmCherry2 and 5 nM linear DNA template (unless indicated otherwise) in a 1.5-mL Eppendorf tube. When expressing proteins from the Min system, the solution was supplemented with 0.5 µM of purified eGFP-MinC, 0.5 µL DnaK Mix (GeneFrontier) and 2.5 mM of ATP. The DNA concentration was adjusted to 6 nM *minD* and 3.8 nM *minE* when both proteins were co-expressed. Next, 5-6 mg of lipid-coated glass beads were added to the swelling solutions. The lipid-coated beads were prepared as described earlier^60^ with minor modifications. The glass beads (0.6 g) were coated with 50 mol % DOPC, 36 mol % DOPE, 12 mol % DOPG, 2 mol % 18:1 cardiolipin, 0.05 mass % DOPE-Cy5 and 1 mass % DSPE-PEG(2000)-biotin with a total mass of lipids of 2 mg. The mixture containing PURE system and lipids was gently rotated at 4 °C for 1 hour to allow for liposome swelling. Four freeze-thaw cycles were applied by dipping the samples in liquid nitrogen for 10 seconds followed by thawing on ice.

### In-liposome gene expression

For liposome samples containing either *YFP*, *btubB*, or *minD* DNA, 0.25 μL of DNase I (NEB, 2 U μL^−1^) was transferred into an empty PCR tube. Using a cut pipette tip, 7 μL of liposomes were transferred to the PCR tubes and mixed with the DNase I by pipetting up and down three times. Liposomes were incubated at 37 °C for 3-16 hours in a thermocycler. The 5.7 × 6.1 mm chambers (length × width) of a µ-Slide 18 Well Glass Bottom (Ibidi) were pre-incubated with 50 µL of 1 mg mL^−1^ BSA in Milli-Q water for 10 minutes and subsequently washed three times with Milli-Q water and once with PB buffer (20 mM HEPES-KOH pH 7.6, 180 mM potassium glutamate, and 14 mM magnesium acetate). Liposomes were diluted 50-fold in PB buffer of which 50 µL was transferred to each chamber. The imaging slide was kept overnight in the dark at 4 °C to allow for the liposomes to sediment.

For experiments with MinD and MinE, the liposome suspension obtained after swelling was diluted 2-fold using a mixture of 1:1 PURE*frex* 2.0 Solution I and Milli-Q water supplemented with 2.5 mM ATP and 0.13 U µL^−1^ DNase I. Next, 10 µL of the diluted liposome sample was transferred to an in-house made, BSA-functionalized round imaging chamber (diameter ∼3 mm). The sample slide was incubated in a laboratory incubator at 37 °C for 2-3 hours before imaging at room temperature.

### Confocal microscopy

Liposomes were imaged using the A1R Laser scanning confocal microscope (Nikon) equipped with an SR Apo TIRF 100× oil immersion objective (NA 1.49). Imaging was performed using laser line 488 nm with emission filter 525/50 nm (gallium arsenide phosphide (GaAsP) detector; YFP and eGFP-MinC), laser line 561 nm with emission filter 595/50 nm (GaAsP detector; PAmCherry2) and laser line 640 nm with emission filter 700/75 nm (photomultiplier tube detector; DOPE-Cy5). For image-based screening, channels 1 (YFP, eGFP-MinC) and 3 (DOPE-Cy5) were imaged simultaneously. Imaging was performed at a fixed dwell time (2.2 µs) and image size (512 × 512 pixels, 0.25 µm pixel^−1^) to optimize for high imaging speed. The pinhole size was 42.1 μm (1 A.U. for 405 nm). Point stimulation was performed using the 405 nm laser for 300 ms at 10-50% laser power (34-167 µW). Image acquisition and stimulation were operated by NIS-Elements version 5.30.07 (Nikon) at room temperature with the focus drift correction module Perfect Focus System (Nikon) enabled.

For optimal stimulation accuracy, point stimulation was performed in the center of the field-of-view (FOV) using the Galvano scanner mirrors. The point stimulation accuracy was regularly checked by the green-to-red photoconvertion of mEos2 using a 3-µL droplet of 50 µM protein in PB buffer sandwiched between a glass slide and coverslip and 1% laser power (5 µW) for 300 ms (**Fig. S8**). When a misalignment of a few pixels was observed, the stage position was adjusted accordingly. The absolute laser powers (in µW) for point stimulation were measured by a Microscope Slide Photodiode Power Sensor with 18 mm × 18 mm Active Area (S170C, THORLABS Germany) placed on top of the 100× objective (without oil) and connected to a Digital Optical Power and Energy Meter (PM100D, THORLABS Germany). The absolute power was measured for 1%, 10%, 25% and 50% relative laser powers while point-stimulating in the center of the FOV. The power value was averaged over 10 s.

Deviations from the above-mentioned protocols for some experiments are described in **Tables S1**, **S7**, **S10** and **S11**.

### Quantification of the lumen intensities of target and off-target liposomes

To quantify the change of PAmCherry2 fluorescence intensity upon stimulation, liposomes expressing YFP were imaged before and 10 seconds after stimulation. Using Fiji (ImageJ2 v2.9.0)^61^, the PAmCherry2 and YFP intensities were measured for the target liposome and all off-target liposomes within a distance of 15 µm. For each liposome, an area was defined manually that covered most of the lumen, of which the average intensity was calculated.

### Training and testing of YOLO

Train and test sets for YOLOv3-tiny^43^ and YOLOv7-tiny^44^ algorithms were generated by manually drawing bounding boxes using the open-source software Label Studio^62^ (https://github.com/heartexlabs/label-studio). While labelling, the individual imaging channels were also inspected using Fiji (ImageJ2 v2.9.0)^61^. The YOLO models were trained and tested on GPU using a Google Colaboratory notebook (available at https://github.com/DanelonLab/Image-based-Phenotypic-Selection/tree/main/Training-YOLO), which was adapted from https://github.com/theAIGuysCode/YOLOv3-Cloud-Tutorial. Unless indicated otherwise, the training of YOLO algorithms was performed from scratch (i.e., without using pre-trained weights) using jpg images of 512 × 512 pixels. During training, saturation and exposure (brightness) augmentations were enabled, but color and rotation augmentations were disabled. The ‘random’ parameter was disabled to prevent up-and downscaling of images during training. For YOLO algorithms trained on 1 class (MinD localized on the membrane), a dataset was used consisting of 700 images containing 688 labelled instances. For YOLO algorithms trained on 2 classes, 40 images were used containing 1,645 instances, 915 without membrane-localized eGFP-MinC (class 1) and 730 instances with membrane-localized eGFP-MinC (class 2). For the 2-class YOLO model (used for detection of dynamical behaviors), special care was taken to exclude (i) small liposomes (approximately <6 µm), which tend to move in and out of focus more frequently than larger liposomes, (ii) liposomes with a blurry, undefined membrane signal, and (iii) liposomes at the edges of the image. The YOLO models were trained and tested using 80% and 20% of the images, respectively. After validation of the 2-class model, a new model was trained on 100% of the data and used for the experiments.

### Real-time image analysis

Real-time image analysis was performed on a HP Z4 workstation with Intel(R) Xeon(R) W-2145 CPU, 64 GB ram, NVIDIA Quadro RTX 4000 GPU and Windows 10 Enterprise operating system. To run YOLO, open-source software CUDA v10.2, cuDNN v8.0.2, OpenCV v4.2.0 and Darknet were installed. Darknet was downloaded from https://github.com/AlexeyAB/darknet and compiled using Visual Studio 2019 as described on https://github.com/AlexeyAB/darknet3requirements-for-windows-linux-and-macos.

Automated image acquisition, image analysis and laser stimulation were controlled by the JOBS module of the NIS-Elements image acquisition software (Nikon). Real-time image analysis by intensity thresholding and simple morphology constraints was performed by the General Analysis module in NIS-Elements (JOB available at https://github.com/DanelonLab/Image-based-Phenotypic-Selection/tree/main/Detection-By-Intensity-Thresholding). This module applied general convolution (size: 3), intensity thresholding, and size (3.5 to 20 µm) and circularity (0.3 to 1) constraints to the YFP channel. For object detection by YOLO, the OutProc function within JOBS exported the image to a local computer folder and initiated a Windows batch script which started YOLO and redirected the resulting coordinates back to NIS-Elements for stimulation (JOB and code available at https://github.com/DanelonLab/Image-based-Phenotypic-Selection/tree/main/Detection-By-YOLO-Single-Time-Frame).

For video-based classification, the liposomes detected in multiple time frames were identified as the same object based on the nearest neighbor with a maximum distance of 5 µm. Bounding boxes with an x-to-y ratio smaller than 0.7 or larger than 1.3 (these were typically liposomes at the edge of the FOV) and/or an area smaller than 37 µm^2^ were excluded from the analysis. Liposomes whose classes changed across the different time frames were classified as dynamic and their coordinates were imported by NIS-Elements for stimulation. The JOB and code for video-based classification are available at https://github.com/DanelonLab/Image-based-Phenotypic-Selection/tree/main/Detection-By-YOLO-Using-Videos. ChatGPT was used to assist in generating parts of the Windows batch scripts.

### Flow cytometry

Liposomes were transferred from the imaging chamber to an Eppendorf tube using a cut pipette tip. The imaging chambers were washed two times with 50 µL of PB buffer, and this volume was added to the same Eppendorf tube. The sample was filtered using the 35 µm nylon mesh of a cell strainer cap of 5-mL round bottom polystyrene test tubes (Falcon). Liposomes were screened and sorted with the BD FACSMelody cell sorter (BD Biosciences) using a 100-µm nozzle, laser line 488 nm with emission filter 527/32 nm (YFP, eGFP-MinC) and laser line 561 nm with emission filter 582/15 nm (PAmCherry2). The photomultiplier tube voltages were set at 342 V for FSC, 405 V for SSC (with SSC threshold 359 V), 646 V for YFP and eGFP-MinC, and 660 V for PAmCherry2. Data were processed with BD FACSChorus version 3 (while sorting) or Cytobank (https://community.cytobank.org/) as shown in **Fig. S9**. The liposomes sorted by FACS were heated at 75 °C for 15 min to inactivate DNase I.

### Quantitative PCR

Quantitative PCR reactions contained 1× PowerUP SYBR Green Master Mix (Applied Biosystems), 400 nM of each primer (1121 ChD and 1122 ChD for *YFP*, 1208 ChD and 1209 ChD for *minD*, 1414 ChD and 1415 ChD for *btubB*) and 1 µL of heat-inactivated liposome sample in a total volume of 10 µL. Quantitative PCR reactions were performed using the Quantstudio 5 Real-Time PCR system (Thermo Fisher). The samples were incubated sequentially for 2 min at 50 °C, 5 min at 94 °C, 45 cycles of 15 s at 94 °C, 15 s at 56 °C, 30 s at 68 °C, followed by 5 min at 68 °C and a melting curve from 65 to 95 °C. The DNA concentration was calculated using calibration curves ranging from 0.00001 pM to 100 pM in the Quantstudio Design and Analysis software v1.4.3 (Thermo Fisher).

### DNA recovery from liposomes and Nanopore sequencing

DNA (*minD*) was recovered from liposomes by PCR. The reactions were set up at 50-µL volume and contained 400 µM of dNTPs, 300 nM of each primer (1525/1526 ChD), 0.4 units of KOD Xtreme Hot Start DNA Polymerase and 2 µL of heat-inactivated liposome sample in 1× KOD Xtreme PCR Buffer. The thermocycling reaction was performed as follows: 94 °C for 2 min, (98 °C for 10 s, 56 °C for 30 s, 68 °C for 84 s) × 39. The amplicons were purified with the QIAquick PCR Cleanup Kit (Qiagen). Recovery of the DNA (1,398 bp) from liposomes was verified using gel electrophoresis and the DNA concentration was measured by the Nanodrop 2000c spectrophotometer (Isogen Life Science). Nanopore sequencing was performed by Plasmidsaurus. Using the Galaxy web platform^63^ (https://usegalaxy.org), the raw sequencing reads were filtered based on size (1,100-1,400 bp) and mapped to the reference sequence using the *Filter FASTQ reads by quality score and length*^64^ and *Map with minimap2*^65^ tools, respectively, using default parameters. The mapped reads were further processed with in-house written R scripts run in Rstudio (R v4.4.1) (https://github.com/DanelonLab/MOSAIC/tree/main/Application-MinD-MTS-Library). The DNA sequences encoding the MTS of MinD were extracted from the mapped reads and subjected to a quality control, whereafter the different DNA variants amongst those sequences were counted.

### Statistics and reproducibility

Error bars in bar plots indicate the standard deviation (SD). The box-whisker plots in **Fig. 2a and Fig. S1** display the median (middle line), the 75/25 percentiles at the boxes and the 90/10 at the whiskers. Statistical tests in **Fig. S1** (Mann-Whitney tests, correlation analysis, linear regression, unpaired t tests) were performed using GraphPad Prism v10.0.3 and were significant if *P* value (two-tailed) < 0.05. Pearson’s *r* is the Pearson correlation coefficient. *R*^2^ is the coefficient of determination. The number of biological replicates (i.e., distinct liposome samples) and technical replicates (repeated measurements) are indicated in the figure legends. The brightness and contrast of microscopy images were adjusted identically for multiple images when comparing fluorescence intensities using Fiji (ImageJ2 v2.9.0)^61^.

## DATA AVAILABILITY

The data are available in the main manuscript and Supplementary Information. Raw and processed data underlying this article, including DNA sequencing data, gel electrophoresis, microscopy images, FACS data, and qPCR data are available in the 4TU.ResearchData repository with DOI: 10.4121/60cdce27-5bd5-44ec-9d8f-53f2ef6b9d58.

**Supplementary Information** includes confocal fluorescence images, FACS plots, qPCR data, performance of object detection models, agarose gel electrophoresis, lists of primers and plasmids, and supplementary tables with experimental details.

## CODE AVAILABILITY

The codes used in this article are available in the 4TU.ResearchData repository with DOI: 10.4121/60cdce27-5bd5-44ec-9d8f-53f2ef6b9d58, as well as on GitHub at https://github.com/DanelonLab under the MIT license, allowing for free use, modification, and distribution.

## ACKNOWLEDGEMENTS

*PAmCherry2.1* gene was a kind gift from the Van der Oost group (Wageningen University, the Netherlands). G571 (pRSET-mEos2) was a kind gift from Kristin Grüßmayer (TU Delft, the Netherlands). We thank Ilja Westerlaken for the purification of PAmCherry2 and mEos2. We thank Rosalie Knot for preliminary experimental work, and Mats van Tongeren for performing preliminary tests with YOLO. We are also thankful to Michal Shemesh, Martin Holub, Timo Kuijt (Nikon Instruments) and the technical support team from Laboratory Imaging s.r.o. for their help with the microscopy automation, Thierry Langlois and Damien Loras from BD Life Sciences – Biosciences for fruitful discussions and testing the BD FACSDiscover S8, and Miao-Ping Chien and the Chien lab for testing their ultrawide-FOV microscope with our liposome samples. MvdB, CD and NJC acknowledge financial support from the “BaSyC – Building a Synthetic Cell” Gravitation grant (024.003.019) of the Netherlands Ministry of Education, Culture and Science (OCW) and the Netherlands Organisation for Scientific Research (NWO). CD acknowledges funding from Agence Nationale de la Recherche (ANR-22-CPJ2-0091-01). NJC acknowledges funding by the NWO-SUMMIT grant “Evolving life from non-life (EVOLF)” (SUMMIT.1.004).

## AUTHOR CONTRIBUTIONS

CD conceived the project; CD and NJC supervised the project and acquired funding; MvdB and CD designed the experiments; MvdB and MS set up the photoactivation pipeline; MvdB performed the experiments and data analysis; MvdB and CD wrote the paper with input from NJC and MS.

## COMPETING INTERESTS

The authors declare no competing interests.

**Correspondence** and requests for materials should be addressed to Christophe Danelon.

## Notes

https://github.com/DanelonLab/Image-based-Phenotypic-Selection

### Competing Interest Statement

The authors have declared no competing interest.

