## Supplementary Information for "Image-based phenotypic sorting of synthetic cells"

**Content:**

Pages S2-10 | Supplementary Figures S1-S9

Pages S11-14 | Supplementary Tables S1-S14

Page S15 | DNA sequences

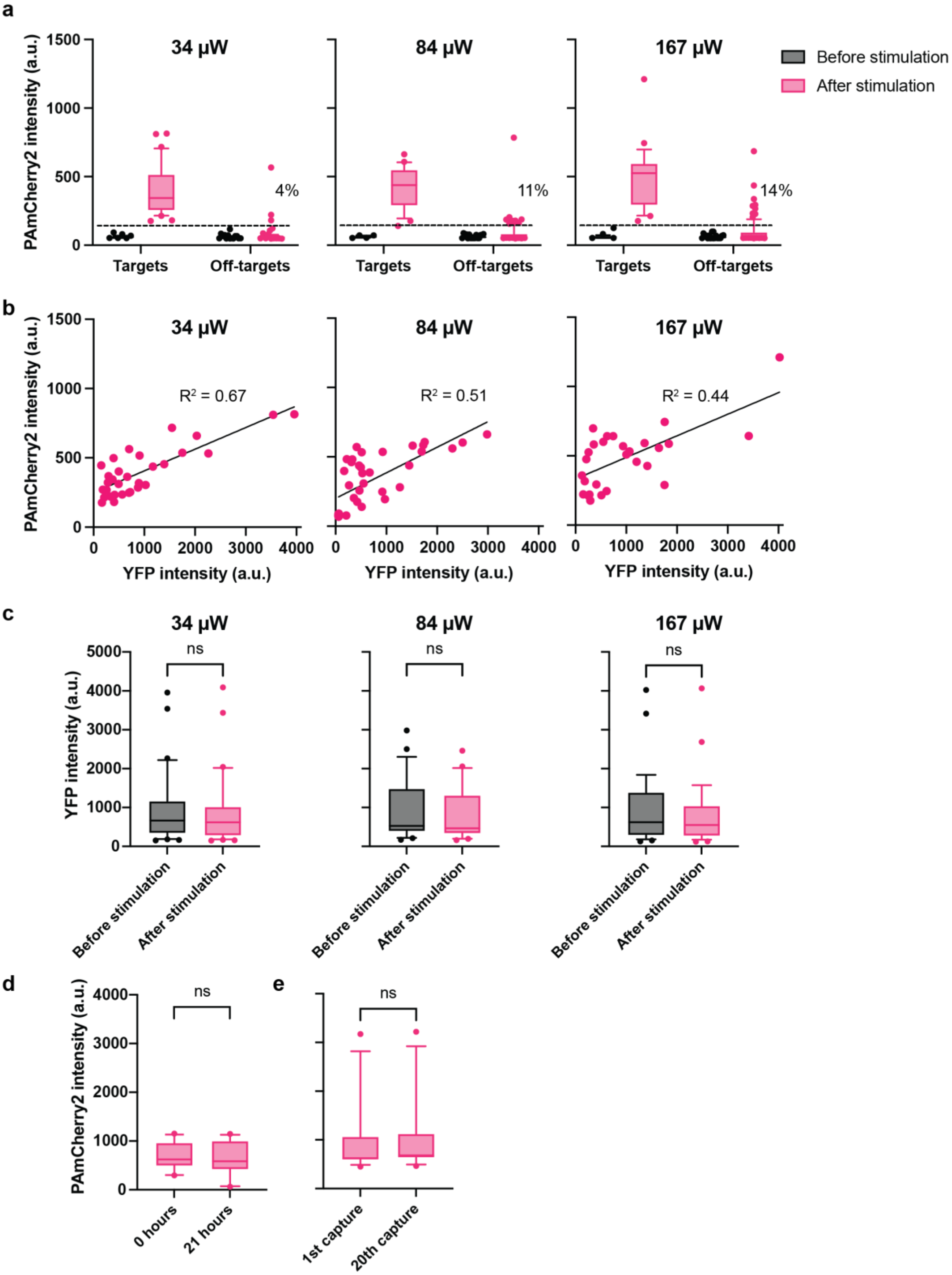


**Fig. S1 PAmCherry2 and YFP fluorescence intensities before and after liposome stimulation.** YFP was expressed in liposomes containing 5 nM *YFP* DNA template and 20 µM purified PAmCherry2. Liposomes were stimulated in the center of the FOV for 300 ms using different laser powers (34, 84 and 167 µW). The fluorescence intensity after stimulation was measured 10 s after stimulation. **a** PAmCherry2 fluorescence intensity in target and off-target liposomes before and after stimulation. All target liposomes were successfully tagged for the three photoactivation conditions with no significant differences in the median intensity values (Mann-Whitney test, $P$ = 0.38 for 34 μW vs 84 μW; $P$ = 0.11 for 34 μW vs 167 μW). All liposomes within 15 μm of the target liposome were considered as off-targets. Per condition, 29-31 target liposomes and 78-94 off-target liposomes were measured. The percentages of off-target liposomes with increased PAmCherry2 intensity (above a certain threshold indicated with the dashed line) relative to all off-target liposomes are appended in the graphs. The data for the target liposomes of the 84 μW condition are also shown in Fig. 2a. **b** Positive correlation between the YFP intensity (before stimulation) and the PAmCherry2 intensity after stimulation (Pearson's $r$ = 0.82 for 34 μW; $r$ = 0.72 for 84 μW; $r$ = 0.67 for 167 μW; $P$ < 0.0001 for all). A linear regression curve is shown in the graphs with the corresponding $R^2$ value. Per condition, 29-31 target liposomes were measured. For the 84 μW condition, ten additional liposomes were measured with low YFP intensity (< 65 a.u.) and low PAmCherry2 intensity (< 95 a.u.) values. **c** YFP intensities before and after stimulation were not significantly different (unpaired t test, $P$ = 0.88, $t$ = 0.1540, degrees of freedom = 60 for 34 μW; $P$ = 0.50, $t$ = 0.6778, degrees of freedom = 56 for 84 μW; $P$ = 0.52, $t$ = 0.6528, degrees of freedom = 56 for 167 μW), thus no significant YFP bleaching occurred. Per condition, 29-31 target liposomes were measured. **d** PAmCherry2 fluorescence values are constant up to 21 hours in liposomes. Twelve YFP-expressing liposomes were activated (34 μW laser power, 300 ms) and tracked over time by imaging every hour. PAmCherry2 intensities at t0 and t21 h were not significantly different (unpaired t test, $P$ = 0.71, $t$ = 0.3782, degrees of freedom = 22). **e** Control experiment for panel d. PAmCherry2 was not bleached due to repeated imaging (20 captures within 30 min) (unpaired t test, $P$ = 0.84, $t$ = 0.2027, degrees of freedom = 22). Twelve target liposomes were measured.

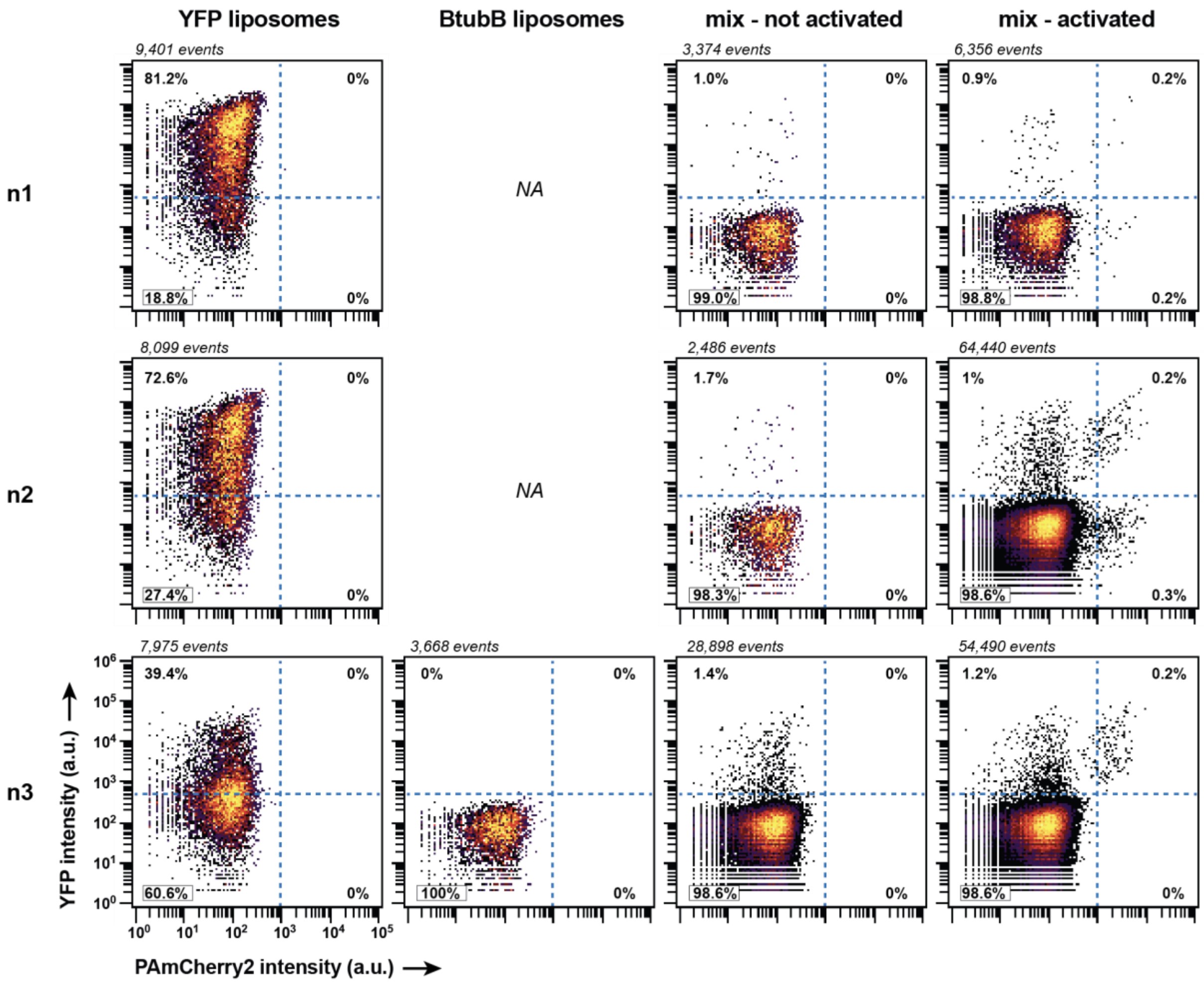


**Fig. S2 Flow cytometry measurements of (a mix of) YFP- and BtubB-expressing liposomes, without or with activation of PAmcherry2.** This figure supplements Fig. 2e with three biological replicates. The two right-most plots from replicate 3 (n3) are also shown in Fig. 2e. The number of events varies per plot as indicated above the graphs. The percentage of liposomes in each gate is also indicated. Experiment 3 (n3) was performed with laser power 34 μW and stimulation in the center of the FOV, while experiments 1 and 2 (n1-2) were performed with laser power 84 μW and stimulation was not restricted to the center of the FOV, likely explaining the lower stimulation precision for experiments 1-2.

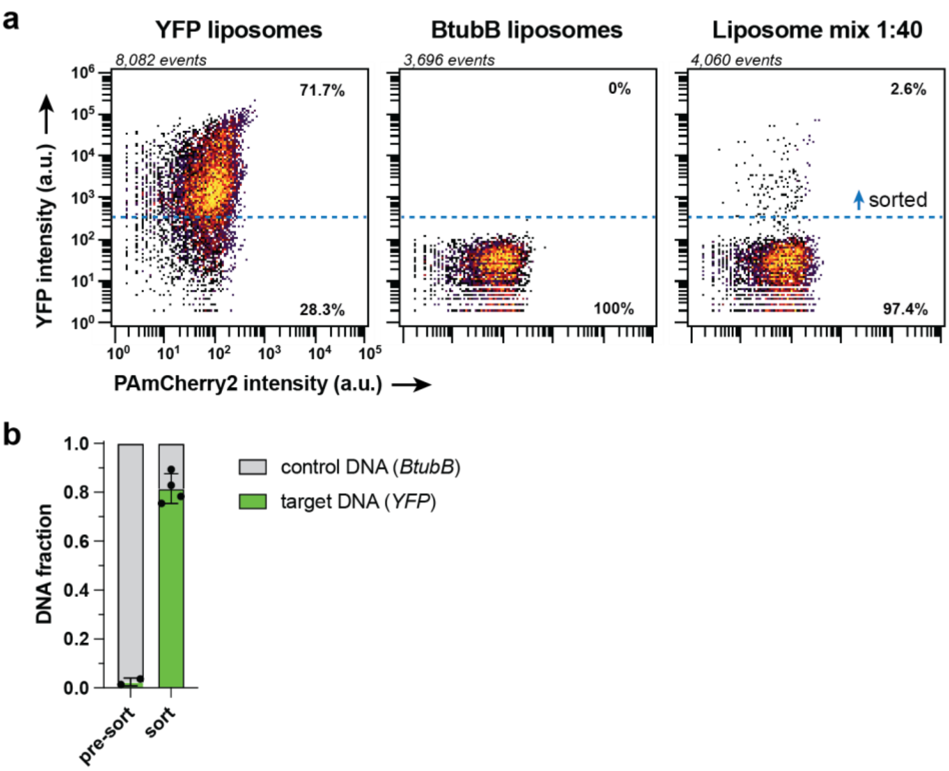


**Fig. S3 FACS sorting of YFP-expressing liposomes from a mixed population of YFP- and BtubB-expressing liposomes based on YFP fluorescence intensity. a** FACS plots showing YFP intensity vs PAmCherry2 intensity for one of two biological replicates. The second replicate corresponds to the left three plots of n3 in Fig. S2. Number of events are indicated above the plots. The percentage of liposomes in each gate is also indicated. **b** qPCR analysis of the sorted liposomes showing enrichment of YFP DNA upon sorting. Two biological replicates (mixed liposome samples) with two technical replicates (sorts) each.

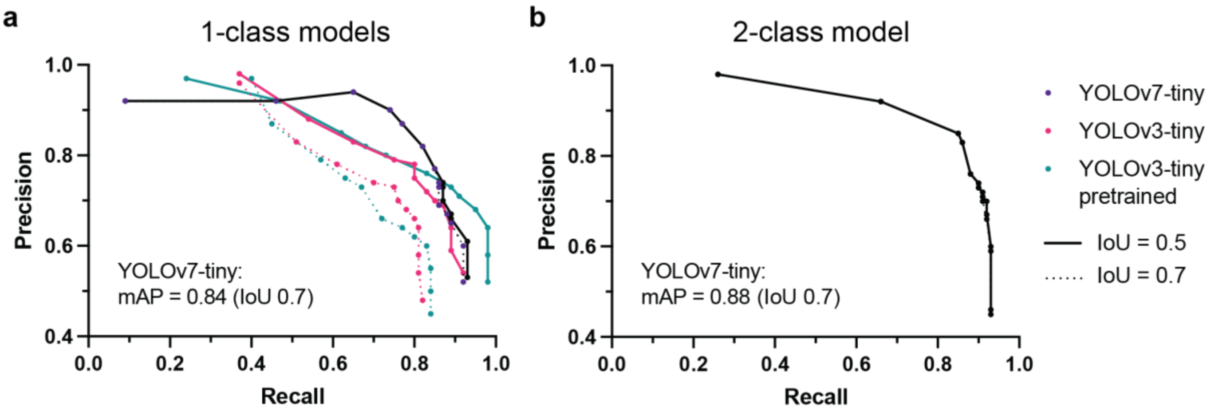


**Fig. S4 Performance of trained YOLO-tiny models. a** YOLO-tiny models trained on one class to detect liposomes with eGFP-MinC at the membrane. mAP is mean Average Precision, and IoU is Intersection over Union. For YOLOv7-tiny (trained from scratch), YOLOv3-tiny (trained from scratch) and YOLOv3-tiny (pretrained), the mAP is 0.85, 0.85 and 0.87 for IoU 0.5, and 0.84, 0.73 and 0.72 for IoU 0.7, respectively. The mAP remains very similar for YOLOv7-tiny when increasing the IoU from 0.5 to 0.7, while for YOLOv3-tiny the mAP decreases. Therefore, the YOLOv7-tiny model has better localization performance of the bounding boxes than the YOLOv3-tiny models. The trained YOLOv7-tiny model was used in liposome experiments at a confidence threshold of 0.8, with both a precision and recall of 82% (IoU 0.5 and 0.7). The YOLOv7-tiny data are also shown in Fig. 3c. **b** YOLOv7-tiny model trained on two classes to detect liposomes with eGFP-MinC at the membrane, and liposomes with eGFP-MinC in the lumen. mAP is 0.88 for both IoU 0.5 and 0.7. The model was used at a threshold of 0.9 (precision 83%, recall 86%).

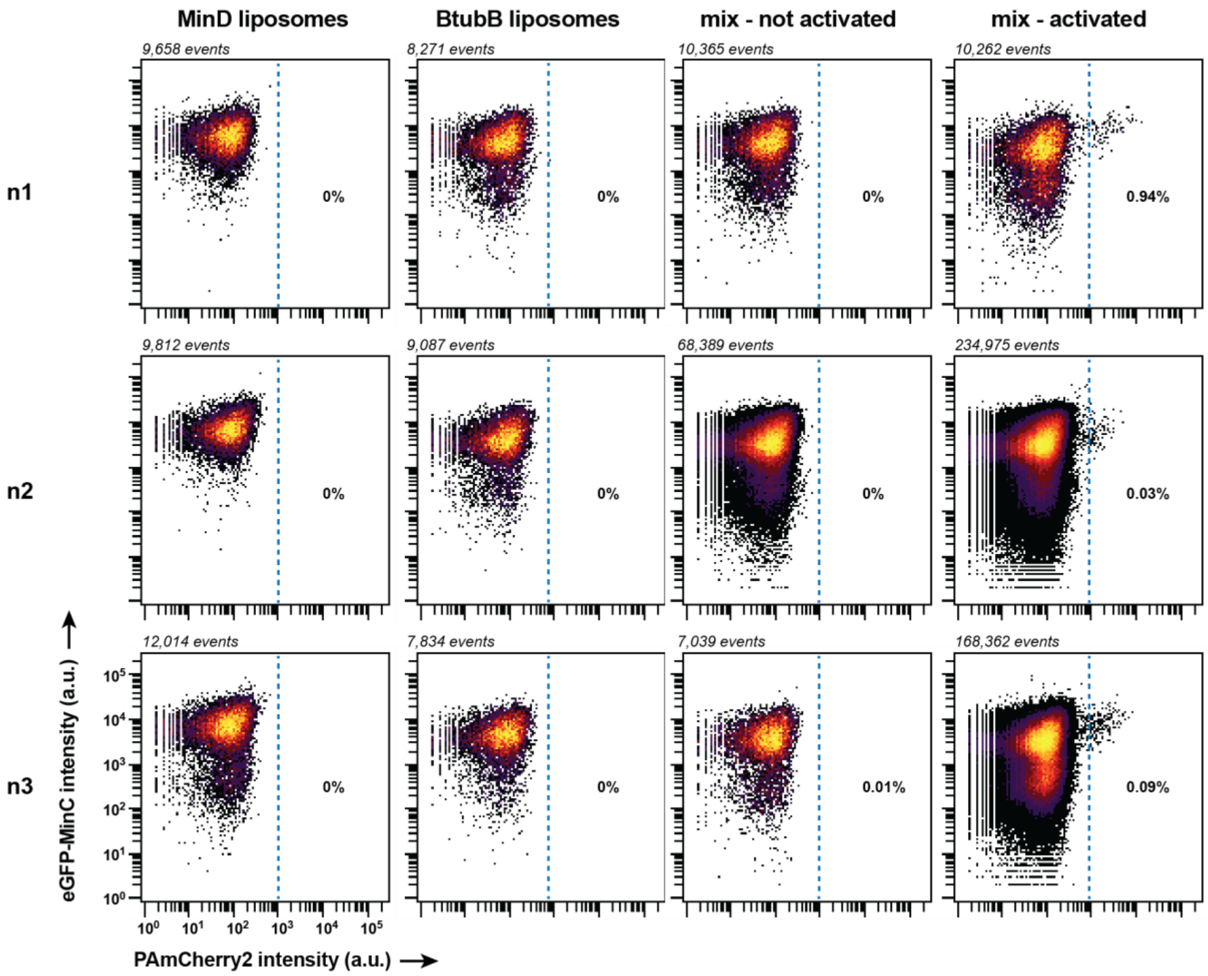


**Fig. S5 Flow cytometry measurements of (a mix of) MinD- and BtubB-expressing liposomes, without or with activation of PAmcherry2.** This figure supplements Fig. 3e with three biological replicates. The two right-most plots from replicate 1 (n1) are also shown in Fig. 3e. The number of events varies per plot as indicated above the graphs. The percentage of liposomes in each gate is also indicated.

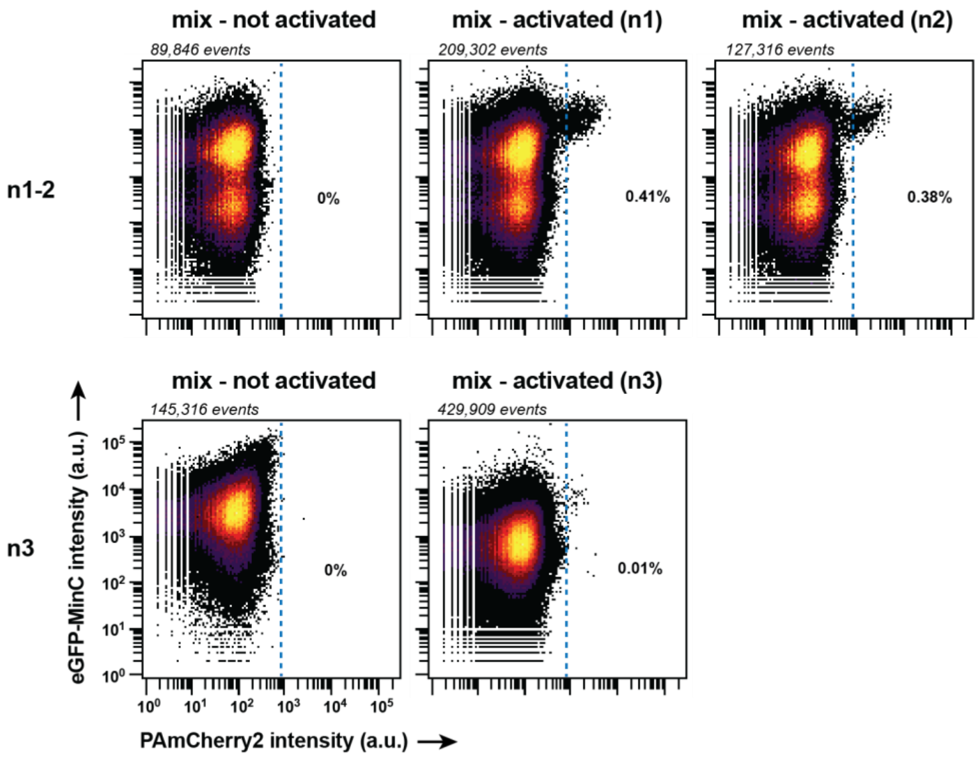


**Fig. S6 Flow cytometry measurements of liposomes expressing a DNA library encoding MinD with mutations in the MTS, without or with activation of PAmcherry2.** This figure supplements Fig. 5. Two biological replicates, for which the first replicate entailed two independent sorts (n1 and n2) and the second replicate entailed one sort (n3). The number of events varies per plot as indicated above the graphs. The percentage of liposomes in each gate is also indicated.

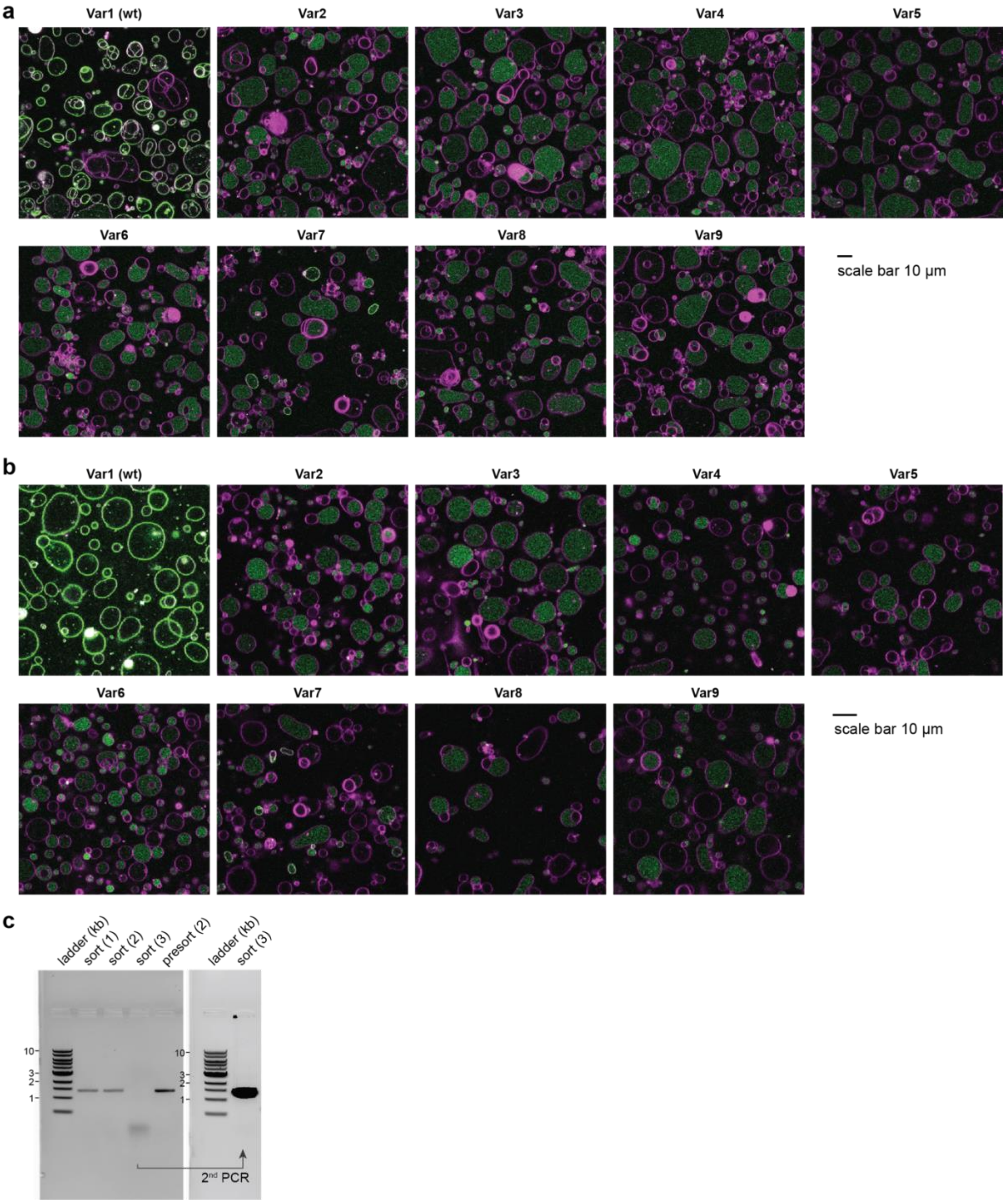


**Fig. S7 Additional data for the library of MinD MTSs. a-b** Fluorescence confocal microscopy images of liposomes expressing clonal DNA variants with mutations in the MinD MTS. Green, eGFP-MinC; magenta, Cy5-conjugated lipids. Scale bars: 10 µm. In replicate 1 (**a**), the liposomes were diluted 15-fold before imaging using PB buffer. In replicate 2 (**b**), the liposomes were diluted 15-fold using a 1:1 mixture of PURE*frex* 2.0 Solution I and MilliQ. **c** Gel electrophoresis with PCR-amplified *minD* (1,398 bp) recovered from sorted liposomes. For sort 3, the full-length DNA was recovered using two subsequent, identical PCRs, where the second PCR contained 1 µL of the first PCR as the source of DNA template instead of the sorted liposomes.

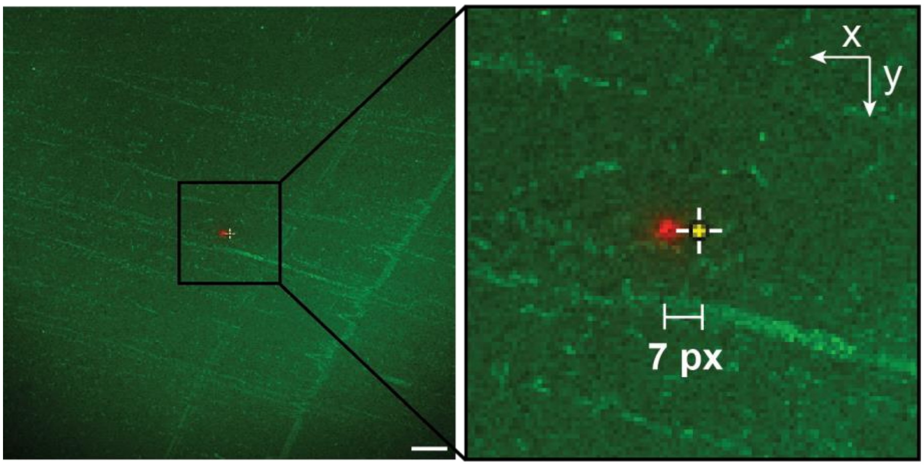


**Fig. S8 Difference between the target and observed location of point-stimulation.** mEos2 protein was imaged at the glass surface of a cover slip and point-stimulated in the center pixel of the field of view (405 nm, 300 ms, 5 µW laser power). The difference in xy location between the target stimulation point (yellow dot) and the observed green-to-red photoconversion and bleaching of mEos2 was quantified (here: 7 pixels in the x-direction, i.e., 1.75 µm). The differences in x and y locations were corrected for in the JOBS protocol (Nis-Elements software). Scale bar: 10 µm.

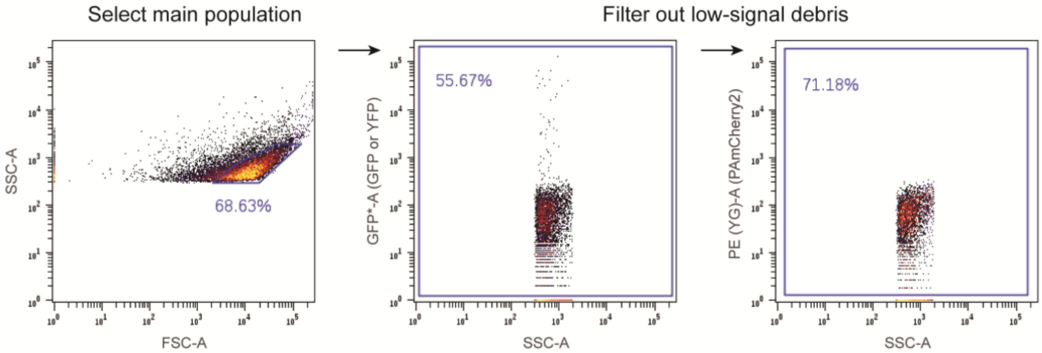


**Fig. S9 Example of gating strategy applied to FACS data.** While sorting liposomes, only the first step (gating based on forward and side scatter light) was applied. Post-measurement analysis involved all three filtering steps.

**Table S1. Experimental details for photoselection and sorting of liposomes from *YFP-btubB* mock library.** The number of stimulated liposomes was estimated by counting the stimulated liposomes for a fraction of all images.

| Exp | Exp ID | Stim at center FOV | Laser power (µW) | Stim time (ms) | Correction xy point stimulation | Number of FOVs screened | Number of stimulated liposomes | FACS sorting threshold | Number of sorted liposomes | Initial ratio *YFP:btubB* |
|---|---|---|---|---|---|---|---|---|---|---|
| 1 | 20230713 | no | 84 | 300 | no | 1,914 | 870 | 1,000 | 193 | 1:40 |
| 2 | 20230721 | no | 84 | 300 | no | 4,687 | 7,218 | 1,800 | 856 | 1:40 |
| 3 | 20240429 | yes | 34 | 300 | yes, 7 px in x direction | 3,440 | 1,668 | 910 | 154 | 1:50 |

**Table S2. Percentages of gated liposomes (extended data for Fig. S2).** Gating thresholds for PAmCherry2 signal and YFP were 988 a.u. and 542 a.u., respectively. ROI 1-4 are the top left, top right, bottom left and bottom right gates, respectively.

| | YFP liposomes | | | | BtubB liposomes | | | | Mix - not activated | | | | Mix - activated | | | |
|---|---|---|---|---|---|---|---|---|---|---|---|---|---|---|---|---|
| Exp | ROI 1 | ROI 2 | ROI 3 | ROI 4 | ROI 1 | ROI 2 | ROI 3 | ROI 4 | ROI 1 | ROI 2 | ROI 3 | ROI 4 | ROI 1 | ROI 2 | ROI 3 | ROI 4 |
| 1 | 81.16% | 0.01% | 18.83% | 0.00% | *NA* | *NA* | *NA* | *NA* | 0.98% | 0.00% | 99.02% | 0.00% | 0.93% | 0.17% | 98.75% | 0.15% |
| 2 | 72.63% | 0.00% | 27.37% | 0.00% | *NA* | *NA* | *NA* | *NA* | 1.73% | 0.00% | 98.27% | 0.00% | 1.02% | 0.18% | 98.55% | 0.25% |
| 3 | 39.39% | 0.00% | 60.61% | 0.00% | 0.00% | 0.00% | 100.00% | 0.00% | 1.38% | 00.0% | 98.62 | 0.00% | 1.21% | 0.22% | 98.57% | 0.00% |

**Table S3. DNA concentrations in the sorted fractions of the *YFP-btubB* mock library as measured by qPCR.**

| | [*btubB*] (pM) | | [*YFP*] (pM) | | *YFP* fraction | |
|---|---|---|---|---|---|---|
| Exp | Pre-sort | Post-sort | Pre-sort | Post-sort | Pre-sort | Post-sort |
| 1 | 0.030018 | 0.000287 | 0.000756 | 0.001045 | 0.0245 | 0.7846 |
| 2 | 0.030467 | 0.000754 | 0.001257 | 0.001159 | 0.0396 | 0.6060 |
| 3 | 0.101736 | 0.000003$ | 0.001403 | 0.000051 | 0.0136 | 0.83507* -0.9525$ |

* when taking the lowest calibration curve data point (0.00001 pM)

$ if calibration curve was forced to extend below the lowest calibration data point (0.00001 pM)

**Table S4. Experimental details for sorting liposomes from *YFP-btubB* mock library based on the YFP signal (liposomes were not stimulated in microscope).** Data consist of 2 biological replicates (mock library liposome mixes), each containing 2 technical sorting replicates.

| Exp | Exp ID | FACS sorting threshold | Number of sorted liposomes | Initial ratio *YFP:btubB* |
|---|---|---|---|---|
| 1 | 20230912 | 350 | 7,021 | 1:40 |
| | | 350 | 5,006 | |
| 2 | 20240429 | 418 | 513 | 1:50 |
| | | 418 | 1,338 | |

**Table S5. Percentages of gated liposomes (extended data for Fig. S3).** Gating thresholds for PAmCherry2 signal and YFP were 988 a.u. and 542 a.u., respectively. ROI 1-4 are the top left, top right, bottom left and bottom right gates, respectively.

| | YFP liposomes | | | | BtubB liposomes | | | | Mix - not activated | | | |
|---|---|---|---|---|---|---|---|---|---|---|---|---|
| Exp | ROI 1 | ROI 2 | ROI 3 | ROI 4 | ROI 1 | ROI 2 | ROI 3 | ROI 4 | ROI 1 | ROI 2 | ROI 3 | ROI 4 |
| 1 | 71.74% | 0.00% | 28.26% | 0.00% | 0.00% | 0.00% | 100.00% | 0.00% | 2.62% | 0.00% | 97.38% | 0.00% |
| 2 | 39.39% | 0.00% | 60.61% | 0.00% | 0.00% | 0.00% | 100.00% | 0.00% | 1.38% | 00.0% | 98.62 | 0.00% |

**Table S6. DNA concentrations in the sorted fractions of the YFP-BtubB mock library as measured by qPCR.** Liposomes were not stimulated in microscope but sorted based on the YFP signal.

| | [*btubB*] (pM) | | [*YFP*] (pM) | | *YFP* fraction | |
|---|---|---|---|---|---|---|
| Exp | Pre-sort | Post-sort | Pre-sort | Post-sort | Pre-sort | Post-sort |
| 1 | 0.190905 | 0.002459 | 0.007178 | 0.008871 | 0.0362 | 0.7829 |
| | | 0.001243 | | 0.010397 | | 0.8932 |
| 2 | 0.101736 | 0.000012 | 0.001403 | 0.000037 | 0.0136 | 0.7542 |
| | | 0.000019 | | 0.000090 | | 0.8289 |

**Table S7. Experimental details for photoselection and sorting of liposomes from *minD-btubB* mock library.** The number of stimulated liposomes for experiment 1 was estimated by counting the stimulated liposomes for a fraction of all images. The exact numbers are presented for experiments 2 and 3.

| Exp | Exp ID | Stim at center FOV | Laser power (µW) | Stim time (ms) | Correction xy point stimulation | Number of FOVs screened | Number of stimulated liposomes | FACS sorting threshold | Number of sorted liposomes | Initial ratio *minD:btubB* |
|---|---|---|---|---|---|---|---|---|---|---|
| 1 | 20231206 | yes | 84 | 300 | Yes, 7 px in x direction | 2,880 | 8,640 | 1,000 | 905 | 1:9 |
| 2 | 20240503 | yes | 34 | 300 | Yes, 7 px in x direction | 5,160 | 1,797 | 700 | 171 | 1:40 |
| 3 | 20240508 | yes | 34 | 300 | Yes, 7 px in x direction | 6,880 | 3,893 | 700 | 189 | 1:40 |

**Table S8. Percentages of gated liposomes (extended data for Fig. S5).** Gate > 988 a.u (PAmCherry2 signal). Percentages based on dataset with most events.

| Exp | MinD liposomes | BtubB liposomes | Mix - not activated | Mix - activated |
|---|---|---|---|---|
| 1 | 0.00% | 0.00% | 0.00% | 0.94% |
| 2 | 0.00% | 0.00% | 0.00% | 0.03% |
| 3 | 0.00% | 0.00% | 0.01% | 0.09% |

**Table S9. DNA concentrations in the sorted fractions of the *minD-btubB* mock library as measured by qPCR.**

| | [*btubB*] (pM) | | [*minD*] (pM) | | *minD* fraction | |
|---|---|---|---|---|---|---|
| Exp | Pre-sort | Post-sort | Pre-sort | Post-sort | Pre-sort | Post-sort |
| 1 | 0.031877 | 0.000262 | 0.005609 | 0.000437 | 0.1496 | 0.6253 |
| 2 | 0.244735 | 0.000030 | 0.007848 | 0.000144 | 0.0311 | 0.8274 |
| 3 | 0.399657 | 0.000005 | 0.016422 | 0.000535 | 0.0395 | 0.9817* |

* when taking the lowest calibration curve data point (0.00001 pM)

**Table S10. Experimental details for photoselection of liposomes displaying Min oscillations.**

| Exp | Exp ID | Stim at center FOV | Laser power (µW) | Stim time (ms) | Correction xy point stimulation | Number of FOVs screened | Number of stimulated liposomes | Time of experiment (h) |
|---|---|---|---|---|---|---|---|---|
| 1 | 20241009 | yes | 34 | 300 | Yes, 11 px in x direction | 170 | 89 | 3.3 |
| 2 | 20241113 | yes | 34 | 300 | Yes, 11 px in x direction | 401 | 190 | 7.8 |
| 3 | 20241203 | yes | 34 | 300 | Yes, 11 px in x direction | 337 | 146 | 7.9 |

**Table S11. Experimental details for photoselection of liposomes from the MinD library with mutations in the MTS.**

| Exp | Exp ID | Stim at center FOV | Laser power (µW) | Stim time (ms) | Correction xy point stimulation | Number of FOVs screened | Number of stimulated liposomes | FACS sorting threshold | Number of sorted liposomes |
|---|---|---|---|---|---|---|---|---|---|
| 1 | 20240522 | yes | 34 | 300 | Yes, 7 px in x direction | 1,720 | 6,155 | 700 | 704 |
| 2 | 20240522 | yes | 34 | 300 | Yes, 7 px in x direction | 1,720 | 5,024 | 700 | 420 |
| 3 | 20240621 | yes | 34 | 300 | Yes, 7 px in x direction | 5,160 | 2,471 | 700 | 88 |

**Table S12. Percentages of gated liposomes (extended data for Fig. S6).** Gate > 988 a.u (PAmCherry2 signal). Percentages based on dataset with most events.

| Exp | Mix - not activated | Mix - activated |
|---|---|---|
| 1 | 0.00% | 0.41% |
| 2 | | 0.38% |
| 3 | 0.00% | 0.01% |

**Table S13. List of primers.** Phosphorothioate bonds are indicated with an asterisk (*).

| Oligo name | Purpose | DNA sequence (5′ – 3′) |
|---|---|---|
| 709 ChD | Gene amplification from T7 promoter to T7 terminator; also used for Sanger sequencing of *minD* variants from G598 containing mutations in the MTS of MinD | CAAAAAACCCCTCAAGACCCGTTTAGAGG |
| 757 ChD | Gene amplification from T7 promoter to T7 terminator | TAATACGACTCACTATAGGG |
| 1429 ChD | *PAmCherry2* gene amplification for the assembly of G595 | AGCGGCCTGGTGCCGCGCGGCAGCCATATGGTGTCTAAAGGGGAAGAGGACAAT |
| 1430 ChD | *PAmCherry2* gene amplification for the assembly of G595 | TTTCGGGCTTTGTTAGCAGCCGGATCCTTACTTATAAAGTTCGTCCATACCCCCCGTTG |
| 1431 ChD | pET15b vector amplification for the assembly of G595 | TAAGGATCCGGCTGCTAACAAAGCC |

| | | |
|---|---|---|
| 1432 ChD | pET15b vector amplification for the assembly of G595 | CATATGGCTGCCGCGCGGCAC |
| 1355 ChD | *mEos2* gene amplification from G571 for the assembly of G577 | AGCAGCGGCCTGGTGCCGCGCGGCAGC CATATGAGTGCGATTAAGCCAGACATG |
| 1356 ChD | *mEos2* gene amplification from G571 for the assembly of G577 | TCCTTTCGGGCTTTGTTAGCAGCCGGATC CTTATCGTCTGGCATTGTCAGGCAATC |
| 1353 ChD | pET15b vector amplification for the assembly of G577 | GGATCCGGCTGCTAACAAAGCCCGAAA G |
| 1354 ChD | pET15b vector amplification for the assembly of G577 | ATGGCTGCCGCGCGGCACCAGGCCGCT G |
| 1121 ChD | DNA quantification (qPCR) *YFP* gene | TGCAACTGGCTGACCACTAC |
| 1122 ChD | DNA quantification (qPCR) *YFP* gene | AATGATTGTCCGGCAGCAGA |
| 1208 ChD | DNA quantification (qPCR) *minD* gene | CGCGACTCTGACCGTATTT |
| 1209 ChD | DNA quantification (qPCR) *minD* gene | AGCATGTCACCTCTGCTTAC |
| 1414 ChD | DNA quantification (qPCR) *btubB* gene | CTGACCCTGCAACGTATTCT |
| 1415 ChD | DNA quantification (qPCR) *btubB* gene | TACGGTTCAGTTTCGCCTTC |
| 1445 ChD | Full-plasmid amplification of G437 to introduce mutations in the MTS of MinD | GCCTTTCTTCTCTTCTTCAATGAAGCGGA A |
| 1448 ChD | Full-plasmid amplification of G437 to introduce mutations in the MTS of MinD | T*T*CCGCTTCATTGAAGAAGAGAAGAAA GGCKWMSWCAAACGCTTGTTCGGAGG ATAAGGATCCGGCTGCTAACAAAGCCC |
| 1525 ChD | Forward PCR primer for amplification of *minD* gene | AAAGTAAGCCCCCACCCTCACATGATAC C |
| 1526 ChD | Reverse PCR primer for amplification of *minD* gene | AAAGTAGGGTACAGCGACAACATACACC ATTTCC |

**Table S14. List of plasmids.**

| Plasmid name | Description |
|---|---|
| G365 | pUC19-*YFP* |
| G376 | pUC57-*minE_opt* |
| G437 | pUC19-*minD* |
| G439 | pUC19-*btubB* |
| G595 | pET15b-PAmCherry2 |
| G571 | pRSET-mEos2 |
| G577 | pET15b-mEos2 |
| G598 | pUC19-*minD* 32-variant MTS library |

### DNA sequences

#### DNA sequence of PAmCherry2

ATGGTGTCTAAAGGGGAAGAGGACAATCTTGCAATCATCAAGGAGTTCATGCGCTTTAAGGTTCAT
CTGGAAGGCTCTGTTAATGGACATGAATTTGAAATCGAAGGAGAAGGGGAGGGTCGTCCTTATGA
GGGAACTCAAACTGCTAAATTGAAGGTAACAAAGGGAGGACCACTTCCGTTCGCGTGGGATATTC
TGTCGCCCCAGTTCATGTACGGAAGCAATGCGTACGTCAAGCATCCCGCTGACATTCCTGACTACT
TTAAGCTTTCCTTCCCCGAAGGTTTCAAATGGGAACGTGTTATGAACTTCGAAGACGGCGGGGTGG
TAACTGTGACACAAGATTCGTCGTTGCAAGACGGTGAGTTTATCTACAAGGTAAAATTGCGCGGGA
CAAACTTTCCCTCTGACGGGCCCGTCATGCAAAAAAAGACCATGGGGTGGGAAACCTTATCAGAG
CGTATGTACCCCGAGGATGGTGCGCTTAAAGGCGAGTTGAAAGCTCGTACTAAGTTAAAGGATGG
CGGACACTATGATACCGAAGTTAAAACGACTTATAAAGCGAAGAAACCCGTGCAATTACCAGGAG
CATACAATGTTAATCGTAAGTTAGACATTACTAGTCACAACGAGGATTATACAATTGTCGAGCAAT
ACGAGCGTGCTGAGGGTCTTCATTCAACGGGGGGTATGGACGAACTTTATAAGTAA

#### Degenerate DNA sequence of 32-variant *minD* library with mutations in the MTS

ATGGCACGCATTATTGTTGTTACTTCGGGCAAAGGGGGTGTTGGTAAGACAACCTCCAGCGCGGC
CATCGCCACTGGTTTGGCCCAGAAGGGAAAGAAAACTGTCGTGATAGATTTTGATATCGGCCTGC
GTAATCTCGACCTGATTATGGGTTGTGAACGCCGGGTCGTTTACGATTTCGTCAACGTCATTCAGG
GCGATGCAACGCTAAATCAGGCGTTAATTAAAGATAAGCGTACTGAAAATCTCTATATTCTGCCGG
CATCGCAAACACGCGATAAAGATGCCCTCACCCGTGAAGGGGTCGCCAAAGTTCTTGATGATCTG
AAAGCGATGGATTTTGAATTTATCGTTTGTGACTCCCCGGCAGGGATTGAAACCGGTGCGTTAATG
GCACTCTATTTTGCAGACGAAGCCATTATTACCACCAACCCGGAAGTCTCCTCAGTACGCGACTCT
GACCGTATTTTAGGCATTCTGGCGTCGAAATCACGCCGCGCAGAAAATGGCGAAGAGCCTATTAA
AGAGCACCTGCTGTTAACGCGCTATAACCCAGGCCGCGTAAGCAGAGGTGACATGCTGAGCATGG
AAGATGTGCTGGAGATCCTGCGCATCAAACTCGTCGGCGTGATCCCAGAGGATCAATCAGTATTG
CGCGCCTCTAACCAGGGTGAACCGGTCATTCTCGACATTAACGCCGATGCGGGTAAAGCCTACGC
AGATACCGTAGAACGTCTGTTGGGAGAAGAACGTCCTTTCCGCTTCATTGAAGAAGAGAAGAAAG
GC**KWMSW**CAAACGCTTGTTCGGAGGATAA

#### DNA sequence of *minE_opt*

TAATACGACTCACTATAGGGGAATTGTGAGCGGATAACAATTCCCCTCTAGAAATAATTTTGTTTAA
CTTTAAGAAGGAGATATACATATGGCGCTGCTGGATTTCTTTCTGAGCCGTAAGAAAAACACCGCG
AACATCGCGAAAGAGCGTCTGCAAATCATTGTTGCGGAGCGTCGTCGTAGCGATGCGGAACCGCA
CTACCTGCCGCAGCTGCGTAAAGATATCCTGGAAGTGATTTGCAAGTATGTTCAAATTGACCCGGA
GATGGTGACCGTTCAGCTGGAACAAAAGGACGGTGATATCAGCATTCTGGAGCTGAACGTTACCC
TGCCGGAAGCGGAGGAACTGAAGTAAGGATCCGGCTGCTAACAAAGCCCGAAAGGAAGCTGAGT
TGGCTGCTGCCACCGCTGAGCAATAACTAGCATAACCCCTTGGGGCCTCTAAACGGGTCTTGAGG
GGTTTTTTG
